# Tissue-resident uterine regulatory T cells support fetal growth

**DOI:** 10.1101/2023.09.07.554921

**Authors:** Laura Maria Florez, Gwladys Fourcade, Guillaume Darrassse-Jèze, Pierre Barennes, Karina Gan-Fernandez, Paul Régnier, Djamel Nehar-Belaid, Tristan Courau, Maria Grazia Ruocco, Serena Zacchigna, Mauro Giacca, Julien Lion, Nuala Mooney, Nicolas Tchitchek, Encarnita Mariotti-Ferrandiz, David Klatzmann

## Abstract

Regulatory T cells (Tregs) are known to contribute to successful allogeneic pregnancy by promoting maternal-fetal tolerance. Here, phenotypic studies and parabionts identified a population of uterine tissue-resident Tregs (uTregs) with trophic functions in the non-pregnant mouse endometrium. Phenotypically, uTregs are CD25^-/lo^ and they present characteristic phenotypic attributes of tissue-resident memory T cells with effector functions. Transcriptionally, uTregs are distinguishable from other peripheral or tissue-resident Tregs by their expression of genes related to extracellular matrix remodeling and vasculogenesis, and they express a polyclonal T cell receptor (TCR) repertoire. Pregnancy triggers the expansion of uTregs, which remain CD25^-/lo^, retain their polyclonal TCR repertoire, and increase the expression of genes involved in regulation of hypoxia and vasculogenesis. Functionally, we show that uTregs support the proliferation of primary uterine microvascular endothelial cells *in vitro* and that a specific depletion of uTregs in pregnant mice leads to fetal intra-uterine growth retardation (IUGR). Thus, a previously unidentified population of CD25^-/lo^ uTregs promotes uterine tissue remodeling before and during pregnancy, and contributes to fetal growth.

## INTRODUCTION

Maternal-fetal tolerance (MFT), the immune tolerance towards an allogeneic fetus during pregnancy, is an essential requirement for mammalian reproduction. The contribution of regulatory T cells (Tregs) to MFT appears crucial as (i) a specific cyclic accumulation of Tregs before fertilization prepares a tolerant environment for a possible embryo implantation (Arruvito et al., 2007; Jasper et al., 2006; Robertson et al., 2009); (ii) during early gestation, self-specific effector memory Tregs expand and are activated in the uterine draining lymph nodes (udLNs) (Chen et al., 2013) and memory Tregs are necessary to establish tolerance in second allo-pregnancies (Rowe et al., 2012); (iii) Treg ablation in mice induces high rejection rates of allogeneic fetuses (Aluvihare et al., 2004; Darrasse-Jèze et al., 2006; Shima et al., 2010; Chen et al., 2013); (iv) women with recurrent spontaneous abortions have low numbers of circulating (Arruvito et al., 2007) and decidual (Inada et al., 2013) Tregs and (v) Treg activation with low-dose IL2 or adoptive transfer of Tregs restores normal pregnancy outcome in a mouse model of immune-mediated spontaneous abortion (Chen et al., 2013; Woidacki et al., 2015).

Although Tregs were detected in several non-lymphoid tissues such as adipose tissue (Feuerer et al., 2009), skin (Seneschal et al., 2012), injured muscle (Burzyn et al., 2013b) and hair follicles (Ali et al., 2017), little is known about Tregs in the uterus. Tissue Tregs differ from Tregs found in secondary lymphoid tissues (SLOs) based on phenotype and function. They are unique for the expression of tissue-specific transcription factors and adhesion molecules, and for their mechanisms of action or expression of effector molecules (Burzyn et al., 2013a).

In humans, Foxp3^+^ Tregs have been observed in uterine biopsies (Inada et al., 2013; Saito et al., 2005, 2007) and fetuses (Darrasse-Jèze et al., 2006), Foxp3 mRNA has been detected in the uterus of healthy women (Jasper et al., 2006), and Tregs have been described to migrate selectively from peripheral blood to the decidua (Kallikourdis and Betz, 2007; Kallikourdis et al., 2007; Tilburgs et al., 2008). In mice, global transcriptome of the uterus revealed the existence of an immune-tolerant environment in which antigen presentation and effector responses are down-modulated after embryo implantation, and which vanishes when Tregs are depleted (Nehar-Belaid et al., 2016). Besides this knowledge, the phenotype, recirculation and function of uterine Tregs are largely unknown.

The role of Tregs in MFT has been associated with thymic-derived Tregs (tTregs) for early pregnancy (Chen et al., 2013) and with both thymic-derived (Rowe et al., 2012) and peripherally converted Tregs (pTregs) for middle and late pregnancy (Rowe et al., 2012, Samstein et al., 2012) in mice. The specificity of these Tregs have been described as towards self-antigens in the early pregnancy (Chen et al., 2013), and later towards fetal-specific antigens (Kahn and Baltimore, 2010; Rowe et al., 2012). Ablation of pTregs induces abortion with signs of abnormal spiral artery (SA) development (Samstein et al., 2012), while it is known that insufficient SA remodeling is linked to preeclampsia, fetal growth restriction, miscarriage, and preterm birth (Ball et al., 2006; Linzke et al., 2014; Lyall, 2002; Pijnenborg et al., 1983, 2006). In this line, uterine NK (uNK) cells have been shown to play a major role in SA development, but not as immune effectors (Croy et al., 2003). uNKs are associated with decidua vascular remodeling and control the depth of invasion of trophoblasts and production of angiogenic factors (Hazan et al., 2010; Smith et al., 2009). The mechanisms by which Tregs could contribute to placental development, directly or indirectly, are unknown. We sought to characterize Tregs in non-pregnant and pregnant uteri and to study their role during early pregnancy. We identified a unique population of tissue-resident memory uterine Tregs (uTregs) under homeostatic conditions. As other tissue-resident Tregs, uTregs have a unique transcriptional profile that appears to be shaped by their tissue environment. They express a plethora of genes involved in extracellular matrix remodeling and blood vessel development, which coordinated expression may be necessary to efficient estrus/menstrual cycle and successful pregnancies. During allogeneic pregnancy, uTregs are activated and proliferate extensively. We show that uTregs favor the growth of primary endothelial cells and that uTregs depletion during early pregnancy leads to markedly reduced fetal growth, a phenomenon known as intra-uterine growth retardation (IUGR), an important cause of fetal and neonatal morbidity and mortality (Sharma et al., 2016). We conclude that, besides the immunological contribution of Tregs to MFT, uTregs have trophic properties ensuring uterine tissue remodeling and likewise fetal growth.

## RESULTS

### The non-pregnant uterine endometrium hosts a unique population of tissue-resident memory CD25^-/low^ Tregs

In order to identify uterine Tregs (uTregs), we used virgin *Foxp3^eGFP^* reporter mice (Wang et al., 2008). Confocal microscopy revealed that uTregs were localized in proximity to the uterine glands (**Figure S1A**). We analyzed by flow cytometry these uTregs based on GFP expression and compared their phenotype with that of Tregs from uterine draining lymph nodes (udLNs). Almost 60% of Tregs of the uterus lacked expression of CD25 compared with only 20% for udLN Tregs (**Figure 1A, 1C** and **Figure S1B**). Tregs represented ∼10% of uterine CD4^+^ T cells, compared to 13% of Tregs amongst udLN CD4^+^ T cells (**Figure 1B**). uTregs expressed levels of CD4 and Foxp3 that were decreased significantly, 1.2- and 1.6-fold, respectively (**Figure S1C** and **S1D**). Moreover, ∼70% of uTregs had a memory phenotype, 44% being effector memory (EM) and 26% being central memory (CM) cells (**Figure S1E**). Remarkably, the CD25^-^ uTregs have mainly an EM phenotype (**Figure S1F**). As the Tregs recovered from uterus could be a mixture of *bona fide* uTregs and circulating passenger Tregs from uterine blood, we performed additional staining on uterine cells harvested after elimination of blood cells through perfusion with PBS immediately before sacrifice. Almost all of the Tregs present in the uterus of blood-depleted mice were CD25^-^ (**Figure 1D**), slightly less than 80% being EM (**Figure 1E**). Thus, most CM, Eff and Nv Tregs observed in the non-perfused uterus (**Figure S1E** and **S1F**) are cells from uterine blood vessels and not from the uterine tissue.

**Figure 1.**
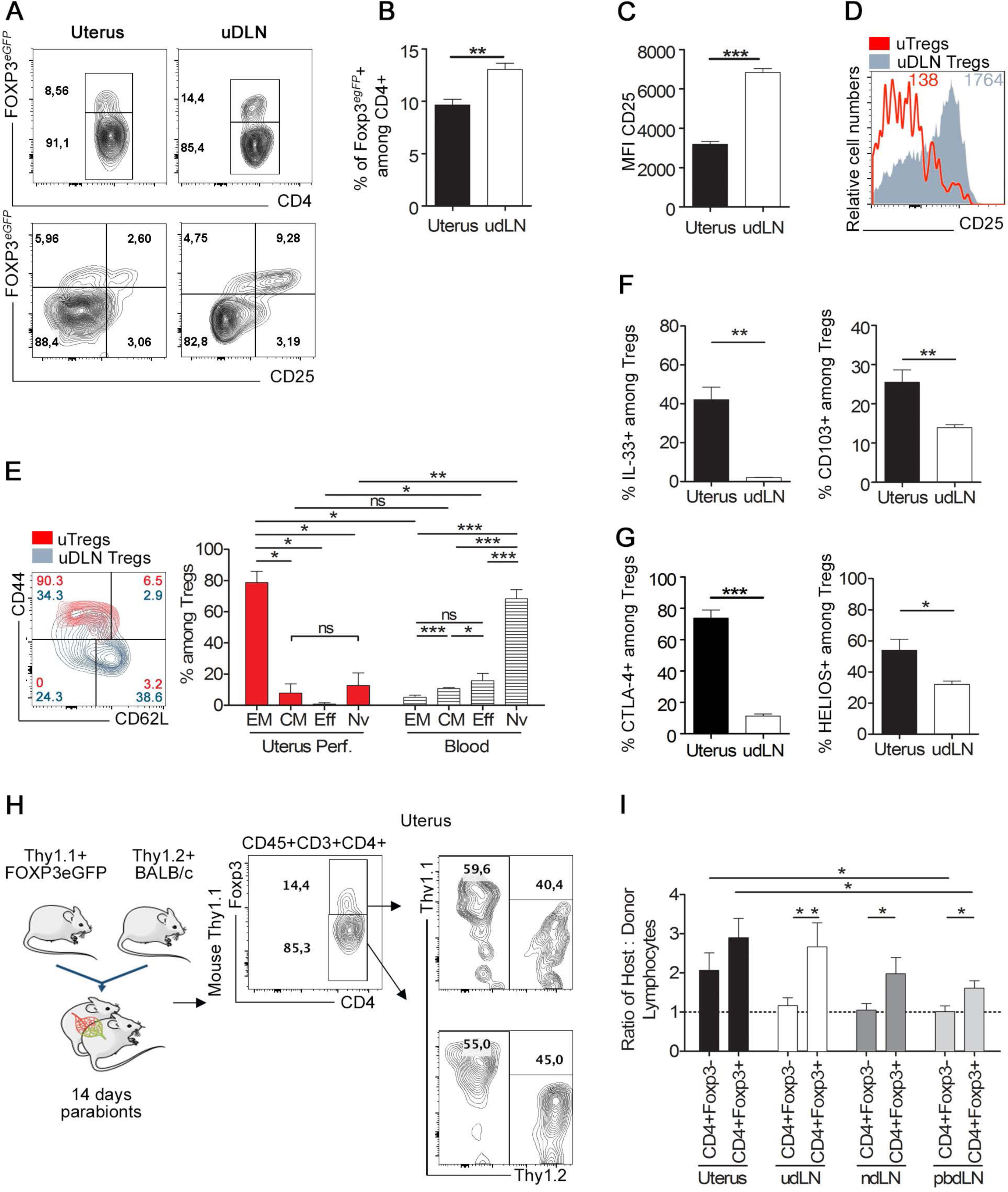
Uterine Tregs are CD25^-^ and have a resident effector memory phenotype with a high proliferation rate. **A)** Gating strategy to identify Treg cells among live CD45^+^ cells in the uterus and udLNs of non-pregnant *Foxp3^eGFP^* reporter mice (C57BL6 background). **B)** Percentages of Foxp3*^eGFP^* -expressing cells among CD4^+^ T cells in the indicated organs (n=12). **C)** Histograms (right) show the expression levels of CD25 as measured by the mean fluorescence intensity (MFI) among Tregs from Uterus and uterus-draining lymph nodes (udLN) (n=12). **D)** Histogram showing CD25 expression in Tregs from perfused uterus (red) and udLN (blue) (right panel). Numbers indicates Mean Fluorescence Index. **E)** Memory populations frequencies among the resident uTregs and blood circulating Tregs. Resident Tregs from perfused uterus are effector memory CD45^+^CD3^+^CD4^+^CD44^+^CD62L^-^CD25^low^Foxp3^+^ cells. (n=4-12 females). (**F-G**): Histograms showing that Tregs from perfused uterus express activation and tissue-resident markers: **F)** Frequency of IL33R and CD103 expressing Tregs. (n=4-12 females). **G)** uTreg expression of CTLA-4 and Helios compared to Tregs from udLNs. (n=4-12 females). **H)** Parabiosis experiment showing the migratory capacity of T cell populations in the uterus and the secondary lymphoid organs. T cells have a high residency in uterine tissue. *Foxp3^eGFPIRES^* BALB/cJ Thy1.1 mice were joined to BALB/cJ Thy1.2. Gating strategy of CD4^+^ T cells in parabiosis: host and donor cells were identified with Thy1.1 (CD90.1) and Thy1.2 (CD90.2) congenic markers among both Foxp3-GFP BALB/cJ Thy1.1 and BALB/cJ Thy1.2 mice. (n=8 parabionts for a total of 16 mice) (*\*p<0.05, **p<0.009, ***p<0.0009*). **I)** The percentages of Thy1.1 and Thy1.2 among CD4^+^Foxp3^-^ and CD4^+^Foxp3^+^ T cells in the uterus, udLNs, parabiosis dLNs (pbdLNs) and non-dLNs (ndLNs) were used to calculate the host-to-donor ratio in each parabiont 14 days after surgery. The dotted horizontal line represents a 1:1 host-to-donor ratio of complete chimerism (n=8 parabionts for a total of 16 mice) (*\*p<0.05, **p<0.009, ***p<0.0009*). EM: effector memory, CM: central memory, Eff: effector, Nv: naïve.

As the phenotype of *bona fide* uTregs is reminiscent of that of tissue-resident effector memory conventional T cells (Trm), we evaluated expression of IL33R and the mucosa integrin CD103, which were previously described as Trm markers. Approximately 40% and 25% of uTregs expressed IL33R and CD103, respectively (**Figure 1F**), while most udLN Tregs did not express these markers. 80% and 55% of Foxp3^+^ uTregs also expressed CTLA-4 and HELIOS, compared to less than 10% and 30% in udLN Tregs, respectively (**Figure 1G**).

To examine the origins, residency and kinetics of replacement of uTregs, we used parabiotic mice generated by surgically joining Thy1.1^+^Foxp3^eGFP^ and wild-type Thy1.2^+^ female mice (**Figure 1H left**). Resident T cells were tracked using the congenic markers Thy1.1 and Thy1.2 (**Figure 1H right**) in the uterus, udLNs, non-draining (ndLNs) and the parabiosis-surgery proximal lymph nodes (pbdLNs). We found that 14 days after parabiosis, peripheral CD4^+^Foxp3^-^ (CD4conv) cells had a host/donor cell ratio close to 1 in SLOs, indicating that full chimerism was reached in these tissues (**Figure 1I**). The host/donor Tregs ratio was the closest to 1 in the pbdLNs, where the highest blood exchange took place. In SLOs, Tregs host/donor cell ratios varied from 1.5 to 3, indicating that Tregs circulate less than CD4conv T cells. Host/donor cell ratios in the uterus varied from 3 to 2 for Tregs and CD4conv subsets, respectively (**Figure 1I**), indicating that the uterine tissue has a lower recirculation and a longer residence time of T cells, which is even more pronounced for Tregs. In line with this and in contrast to udLN Tregs, uTregs are significantly enriched for EM cells negative for CCR7, the lymph node homing receptor (**Figure S1G-S1I**).

Finally, Tregs sorted from the uterus of non-pregnant mice appeared as potent as splenic Tregs at suppressing the proliferation of CD4^+^Foxp3^-^ splenic Tconv cells (**Figure S1J**).

Thus, a previously uncharacterized population of CD4^low^Foxp3^low^CCR7^-^CD25^-^CTLA-4^high^ Tregs is present in the virgin uterine endometrium and their phenotype and residency qualify them for being resident effector memory uTregs.

### The uterine microenvironment fosters uTregs cycling in the absence of IL-2

We next evaluated the homeostatic cycling status of uTregs. We observed that at steady state a high proportion (∼70%) of uTregs expressed the proliferation marker Ki67, far higher than the ∼10% for Ki67^+^ Tregs in udLNs (**Figure 2A**). To assess whether Ki67 expression revealed uTregs division or arrest in G1 (Bullwinkel et al., 2006), we treated mice with the EdU nucleoside analogue and analyzed EdU incorporation into replicating DNA of dividing cells 16 hours later. We found on average that after a single EdU injection, 24.5% of Tregs in the uterus were EdU^+^, compared to 3.8% in udLNs, 5.8% in ndLNs and 4.5% in spleen (**Figure 2B**).

**Figure 2.**
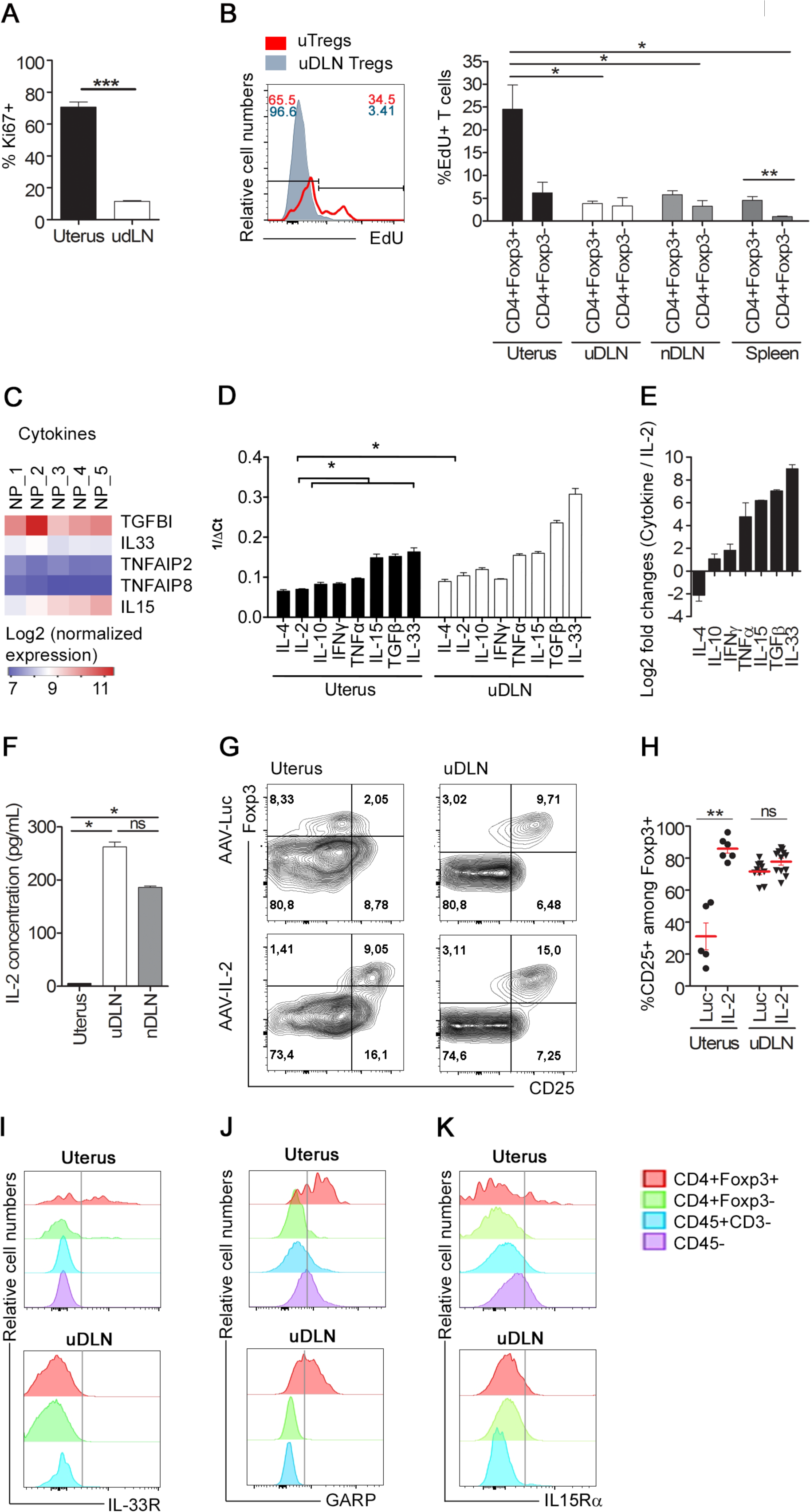
Low CD25 expression in uTregs is a result of an IL2-deprived environment. (**A-B**) uTregs proliferate: **A)** 70% of uTregs are Ki67^+^, percentages of Ki67^+^ Foxp3^+^ cells in non-perfused (n=5, left panel) and perfused (n=4) uterus and udLNs (right panel) from C57BL6 mice. **B)** Histogram representing Edu-labeled cells among uTregs and udLN Tregs (left panel). Percentages of Edu^+^ Tregs and CD4conv cells in the uterus, udLNs, ndLNs and spleen of non-pregnant animals (n=4, *p<0.05). **C)** Log2 of quantile normalized expression for all the genes belonging to the interleukin, tumor necrosis factor (TNF) and interferon families and their corresponding receptors from the HGCN database within the GSE68454 dataset. IL15, TGF-β, IL33 and TNF are present in the non-pregnant uterus. **D)** Relative expression levels (1 / ΔCt) of T cells related to cytokines in the uterus and udLNs. Differences among the highly expressed cytokines—IL15 TGF-β and IL33—and poorly expressed cytokines—IL-4, IL2, IL-10, IFNɣ and TNFα—are significant in the uterus (*p=0.0286*) and udLNs (*p=0.0079*). IL2 levels are lower in uterine tissue than in lymphoid tissue (*p=0.0159*) (n=4). (**E-F**) IL2 is scarce in the non-pregnant uterus: **E)** Log2 fold change in the relative expression of uterine cytokines vs. IL2 by qPCR (*p=0.0286*) (n=4). **F)** Cytokine IL2 concentration measured by ELISA IL2 from the non-pregnant uterus, udLNs and ndLNs (n=4). **G)** Uterine Treg cells in C57BL/6 mice infected with AAV-IL2 regain CD25 expression. Dot plots of Foxp3 and CD25 among CD4^+^ cells in the uterus and udLNs after AAV-IL2 treatment compared to the AAV-Luc control. **H)** Percentages of CD25^+^ cells among Tregs (*p=0.0043*) after AAV-IL2 treatment (n=6 representative of 2 independent experiments). **I)** IL33R, **J)** GARP and **K)** IL15Rα are expressed by uTregs. Histograms showing IL33R^+^, GARP^+^ and IL15Rα^+^ cells among CD45^-^ (violet), CD45^+^CD3^-^ (blue), CD3^+^Foxp3^-^ (green), Foxp3^+^ (red) uterine (upper panels) and udLN (lower panels) populations from C57BL/6 mice.

Cytokines have an important role in Tregs homeostasis and division. We used transcriptomic data from total uterine tissues (Nehar-Belaid et al., 2016) to analyze the cytokines present in the uterine microenvironment that could support resident uTregs survival and cycling. We could not detect IL-2, but found that the non-pregnant uterus expressed transforming growth factor β (TGFβ), interleukin 15 (IL-15), interleukin 33 (IL-33) and tumor necrosis factor α (TNFα) (**Figure 2C**). We validated these results by quantitative RT-PCR (qPCR) with uterine tissue from an independent experiment. We found that IL-33, TGFβ, IL-15 and TNFα were the most highly expressed cytokine RNAs in the uterine tissue. In contrast, IL-4, IFNγ, IL-10 and IL-2 RNAs were expressed at low levels, with IL-2 RNA levels being 21-fold lower in uterine tissues than in udLNs (**Figure 2D**). Compared to IL-2 RNA expression, IL-15, TGF-β or IL-33 levels were respectively 74-, 133- and 508-fold higher (**Figure 2E**). We confirmed by ELISA that no IL-2 protein expression could be detected in the uterus, compared to 200-300 pg/mL in udLNs and ndLNs (**Figure 2F**).

We then investigated whether the absence of IL-2Rα (CD25) expression on uTregs was directly related to the low level of IL-2 in the uterine microenvironment by assessing the effect of exogenous IL-2 on CD25 expression by uTregs. Mice received a single injection of a recombinant adeno-associated virus (rAAV) mediating constitutive expression of IL-2 (AAV-IL2) or of luciferase as a control (AAV-LUC) (Churlaud et al., 2014). Fourteen days later, overexpression of IL-2 had turned most of the CD25^-^ uTregs into CD25^+^ cells (**Figure 2G-H**). Thus, the lack of CD25 expression by uTregs is a consequence of the absence of IL-2 in the uterus.

IL-33, TGF-β and IL-15 are the most expressed cytokines in the uterine tissue. As these three cytokines are functionally relevant to Tregs maintenance, differentiation and proliferation, we investigated whether their receptors were expressed by uTregs. Approximately 40% of uTregs express IL-33R, while no expression of this receptor was found in udLN Tregs or other cell populations in the uterus (**Figures 1F and 2I**). ∼60% of uTregs expressed the receptor for latent TGF-β (GARP), as did udLN Tregs (**Figure 2J**). ∼20% of uTregs expressed IL-15Rα (**Figure 2K**), while no expression of IL-15Rα could be detected on Tregs or CD45^+^ cells in udLNs (**Figure 2K**); moreover, 7% of CD3^+^ and 34% of CD45^-^ cells from the uterus expressed IL-15Rα (**Figure 2K**), which can be used for IL-15 trans-presentation (Dubois et al., 2002). Altogether, the uterine milieu is rich in IL-33, TGF-β and IL-15, which could replace IL-2 in uTregs maintenance (Lodolce et al., 1998).

### uTregs from non-pregnant mice express a unique transcriptional program with enhanced expression of genes involved in extracellular matrix remodeling and vasculogenesis

We compared the transcriptomes of purified uTregs to that of CD4^+^Foxp3^-^ uterine helper T cells (uCD4conv) and of Tregs from udLN, all from non-pregnant mice. Principal component analysis (PCA) of the samples’ global transcriptomes clearly distinguished uTreg cells from uCD4conv cells, but also from udLN Tregs (**Figure 3A**). This indicates that uTregs have a unique Tregs transcriptomic signature that distinguishes them from LNs Tregs, as previously described for other tissue Tregs (Feuerer et al., 2009; Cipolletta et al., 2012; Burzyn et al., 2013b; Kolodin et al., 2015; Arpaia et al., 2015; Vasanthakumar et al., 2015; Kuswanto et al., 2016; Ali et al., 2017).

**Figure 3.**
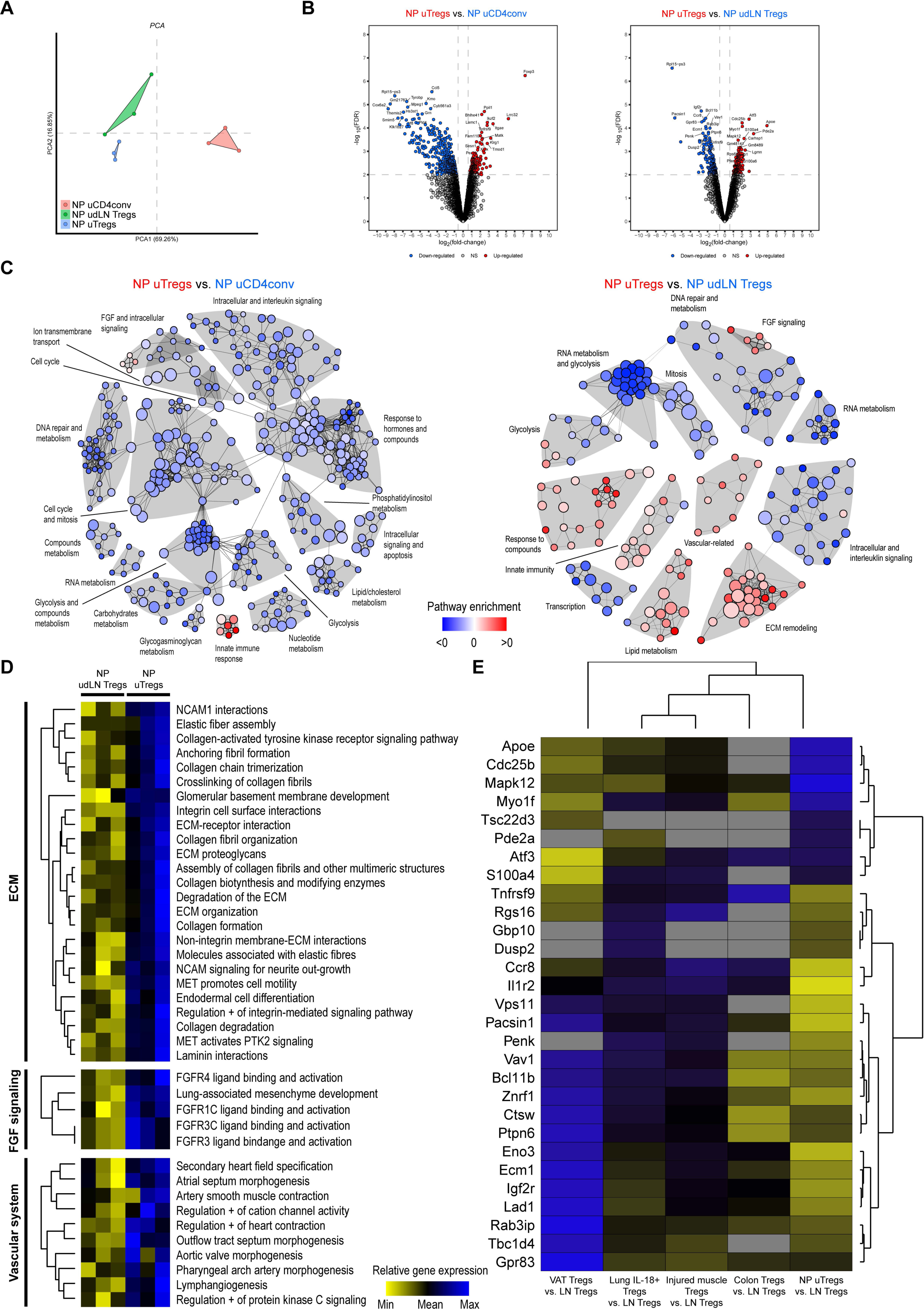
Unique uTregs transcriptomic program highlights a specific extracellular matrix remodeling function. RNA-Seq analysis from purified CD25-uTregs, uterine CD4^+^Foxp3^-^ (CD4conv) T cells and udLN CD25^+^ Tregs (n=3). **A**) PCA of transcriptional profiles distinguishing NP uTregs, NP uterine CD4conv and NP udLN Tregs. **B)** Volcano plot showing genes differentially expressed between NP uTregs and NP uterine CD4conv (left) and between NP uTregs and NP udLN Tregs (right). **C**) Network representation of pathway enrichment analysis from the list of pathways statistically differentially expressed between NP uTregs and the indicated groups. Each node in the representation corresponds to a pathway, and nodes are connected if the lists of genes in the associated pathways are overlapping. Nodes are colored using a color-gradient scale ranging from blue (<0) to red (>0) according to the difference of mean expressions between the conditions. Clusters of pathways are distinguished from each other by their grey-colored area. **D**) Details of the enrichment of biological pathways related to observed ECM, FGF signaling, and vascular system clusters in the enrichment network shown in the NP uTregs vs. NP udLN Tregs comparison. Expression score for each pathway found to be statistically differently expressed in the different individual are represented using a color-gradient scale ranging from yellow (low) to blue (high). **E**) Heatmap of comparison between differentially expressed genes vs. bulk lymph-nodes cells (LN) of Tregs from NP or E6-E8 uterus, and Tregs from visceral adipose tissue (VAT), injured muscle, colonic lamina propria and lung IL18^+^ Tregs. Blue boxes indicate up-regulation of the gene in the given condition, yellow boxes indicate down-regulation. Grey boxes indicate NA values.

The analysis of the top differentially expressed genes (DEGs) between uTregs and uCD4conv showed that uTregs expressed genes of a classical Tregs signature (*Foxp3, Helios/Ikzf2*), with an activated phenotype (*CD103/Itgae, Klrg1, GARP/Lrrc32*), as well as many genes related to tissue-residency, tissue homing, cell-to-matrix adhesion and angiogenesis (**Figure 3B left**). Top DEGs between uTregs and udLN Tregs (**Figure 3B right**) include genes involved in activation/proliferation and genes involved in cell-matrix interactions, tissue repair and angio/vasculogenesis. Of note, among the latter ones, *Atf3* is a crucial factor for vascular regeneration (McDonald et al., 2018) and is a negative regulator of inflammation in human fetal membranes (Lim et al., 2016). Compared to udLN Tregs, uTregs over-express pathways linked to Extracellular matrix (ECM) remodeling, vascular system, and FGF signaling (**Figure 3C** and **3D**).

Noteworthily, uTregs transcriptome differs from that of previously described tissue-resident muscle, VAT, colon, and lung Tregs (**Figure 3E**). Thus, the uTregs unique transcriptome appears to be influenced by their environment as it is the case for other tissue resident Tregs. They differ from circulating Tregs and from Tregs from other tissues by increased expression of genes related to ECM remodeling and vasculogenesis, suggesting a role of uTregs in the remodeling of the uterus in the menstrual cycle.

### uTregs are activated and accumulate during pregnancy

Early pregnancy (E8) resulted in a 4-fold increase of uTregs number compared to that of the non-pregnant (NP) uterus (**Figure 4A**). Pregnancy induced an increase in the proportion of EM and Eff uTregs and a decrease of CM and naïve uTregs (**Figure 4B**). In contrast, there were no significant changes in the functional phenotype of Tregs from the udLNs, except for the Eff fraction, which increased at E8 (**Figure S2A**). CD25 expression on uTregs remained remarkably low during pregnancy, and even decreased at E12, in contrast to its increase on udLN Tregs (**Figure 4C**), perhaps reflecting the lower IL-2 expression in the pregnant vs. virgin uteri (**Figure S2B**). Around 40% of total uTregs maintained expression of CD103 (**Figure 4D**), suggesting that the resident phenotype of these cells is conserved during early pregnancy. Pregnancy triggered an increase in CTLA-4 and Ki67 expression that reached approximately 90% and 80% (**Figure 4E and 4F**) of total uTregs, respectively, while HELIOS remained stably expressed in 60% of uTregs (**Figure 4G**).

**Figure 4.**
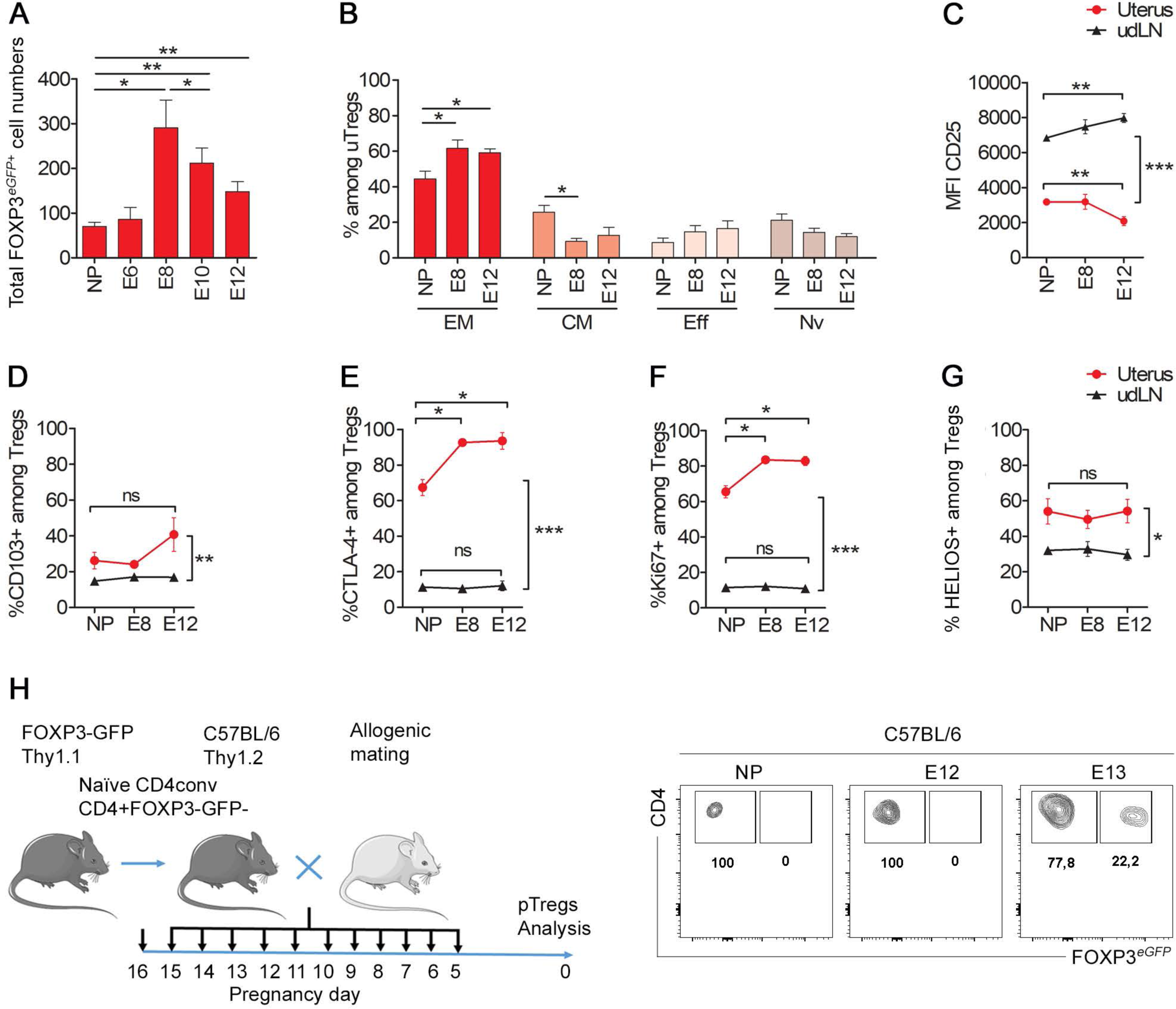
uTregs expand during pregnancy and retain their uTreg phenotype. **A)** uTregs cell numbers increase during the allogeneic pregnancy of female Foxp3*^eGFP^* (C57BL/6) mice mated with BALB/C males. Histograms show the numbers of Tregs (*p<0.01*) among total uterine cells. **B)** Tregs present an activated memory phenotype at E8 and E12 days of pregnancy. Percentages of memory populations among uTregs. (n=3 to 5). **C)** CD25 is modulated among Tregs during pregnancy. Line bar graphs show the expression levels in MFI of CD25 in Tregs, uterus (red line) and udLN (black line) tissues. Data from NP, E8, and E12 of pregnancy. (*\*p<0.05, **p<0.01, ***p<0.001*) (n= 3 to 16 mice representative of 2 different experiments). **(D-G):** Percentages of **D)** CD103^+^ **E)** CTLA-4^+^, **F)** Ki67^+^ and **G)** Helios^+^ cells among uterine or udLN Tregs. Data from NP, E8 and E12 uterus (red line) and udLN (black line) tissues (*\*p<0.05, **p<0.01, ***p<0.001*) (n= 3 to 16 mice representative of 2 different experiments). **H)** Adoptive transfer of Thy1.1 CD4^+^GFP^-^ cells into C57BL/6 females BALB/c mated. Gating strategy showing the induction of uTregs determined by the expression of Foxp3-GFP. pTregs appear at E13 of mid-gestation. Data representative of 3 independent experiments.

As inducible Tregs (pTregs) have been shown to play a role in pregnancy (Samstein et al., 2012), we assessed whether we could detect them in the uterus of pregnant mice. We performed adoptive transfer of sorted Thy1.1^+^CD4^+^GFP^-^CD62L^+^CD44^-^ naïve T cells harvested from *Foxp3^eGFP^* females into Thy1.2^+^C57BL/6 females which were then mated with BALB/c male. We could detect few CD4^+^Foxp3GFP^+^Thy1.1^+^ pTregs in the pregnant uteri, but only from E13 (**Figure 4H**). These results indicate that the uTregs accumulating during early gestation are exclusively of thymic origin.

### uTregs have a polyclonal repertoire

Tissue-specific Tregs were shown to express tissue-specific repertoires (Feuerer et al., 2009; Burzyn et al., 2013b). We sought to evaluate the T cell receptor (TCR) repertoires of uTregs and udLN Tregs before and during pregnancy (**Figure S3**). We observed that the TCR repertoire of NP uTregs is polyclonal, with a usage of variable genes similar to that of Tregs from the udLN and the uterus of pregnant animals. Compared to TCR repertoires from control tissues, the NP uTregs repertoires contain slightly more clonotypes that are detected more than 20 times (**Figure 5A**). Third, the proportion of such expanded clonotypes is almost 6-fold higher in the uTregs from pregnant mice (E6-E8 uTregs) than in NP uTregs (approximately 60% vs 10%, respectively). Thus, in the pregnant uterus, uTregs contain multiple expanded clonotypes, in line with their high Ki67 index (**Figure 2A, 2B and 4F**). To get more insights in the specificity of uTregs, we evaluated clonotype sharing between the repertoires of uTregs from non-pregnant and pregnant uteri (NP uTregs and E6-E8 uTregs) and that of control tissues (**Figure 5B**). We found that the sharing of clonotypes between the NP uTregs and control Tregs repertoires is around 1%, while its sharing with E6-E8 uTregs repertoires is modestly increased to around 3% (**Figure 5B**). Compared to the sharing of NP uTreg repertoires, the sharing of the E6-E8 uTregs repertoires with control repertoires as well as with NP uTregs repertoires is uniformly increased by almost 5-fold. This indicates that the clonotypes that expanded in E6-E8 uTregs are more public than those present in NP uTregs repertoires. Of note, 30% of the E6-E8 uTregs TCRs that are shared with NP uTregs TCRs are also shared with TCRs from other tissues (**Figure 5C**). Finally, shared TCRs have a higher probability of generation than private TCRs (**Figure 5D**) in agreement with the literature (Quigley et al., 2010; Madi et al., 2014). Altogether, the polyclonality of the uTreg repertoires and the sharing with TCRs from unrelated tissues indicate that the TCRs of uTreg are mostly not specific for uterine-specific antigens.

**Figure 5.**
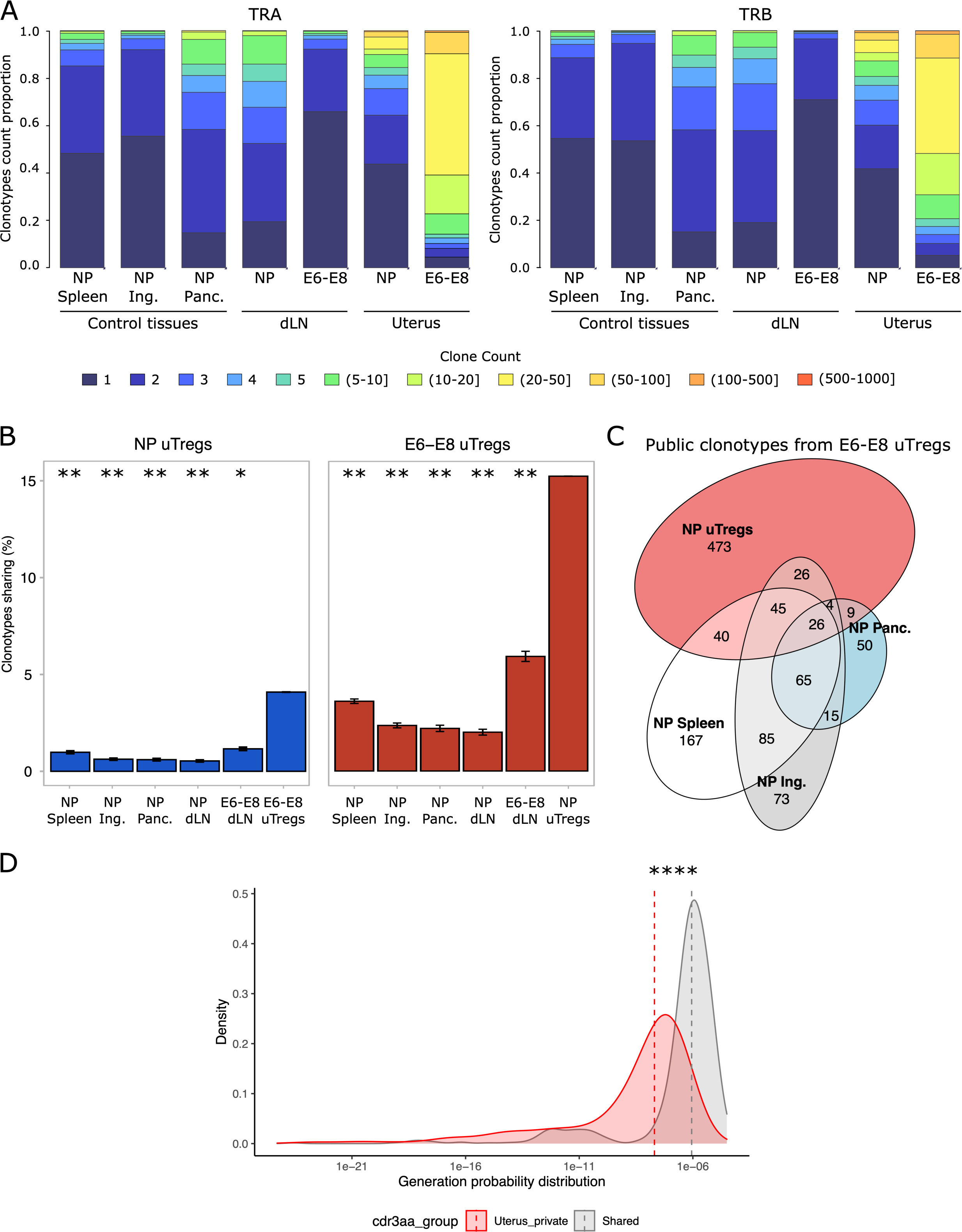
TCR repertoire of uTregs. **A)** Distributions of TCR sequence occurrences for each repertoire. TCRα (left) and TCRβ (right) clonotypes count proportion of Tregs from different organs – spleen (NP Spleen), inguinal (NP Ing.) and pancreatic (NP Panc.) non draining lymph nodes (ndLN) as well as uterine draining lymph node (udLN) and uterus from non-pregnant (NP) and pregnant (E6-E8) mice. Clonotypes are classified according to their occurrence whose proportion/distribution is represented as a cumulative histogram. Each color corresponds to counts as indicated in the legend. **B)** Clonotypes sharing comparison. Proportion of clonotypes shared between compared sample (label on bottom of each barplot) for NP uTregs and E6-E8 uTregs samples. To normalize, sub-sampling was performed according to the smallest sample for each comparison (15 594 for the E6-E8 uTregs analysis and 4 166 for the NP uTregs analysis). T-test was performed using uterine Tregs as control for each analysis: P > 0.05 is NS; *P ≤ 0.05; **P ≤ 0.01; ****P ≤ 0.0001. **C)** E6-E8 uTregs public clonotypes sharing. The Euler Venn diagram represents the public clonotypes of E6-E8 uTregs shared with control tissues (NP Spleen, NP Inguinal, NP Pancreatic) and NP uTregs. **D)** Probability of generation of shared and private uTregs clonotypes. For each TCRβ CDR3aa from E6-E8 uTregs, the generation probability distribution was compared between public CDR3aa in grey (shared with at least one of the controls tissues, NP Spleen and/or NP Inguinal and/or NP Pancreatic) and “uterus specific” CDR3aa in red (E6-E8 uTregs privates or shared with NP uTregs). T-test was performed: P > 0.05 is NS; *P ≤ 0.05; **P ≤ 0.01; ****P ≤ 0.0001.

### Gene expression profiling of uTregs from pregnant mice highlights trophic functions and cell proliferation

We compared the whole transcriptome of uTregs from E6 to E8 of allogeneic pregnancy (E6E8 uTregs) to that of uCD4conv T cells (NP uCD4conv), udLN CD25^+^ Tregs (NP udLN Tregs) and resident uTregs (NP uTregs) from the non-pregnant mice (**Figure 6**).

**Figure 6.**
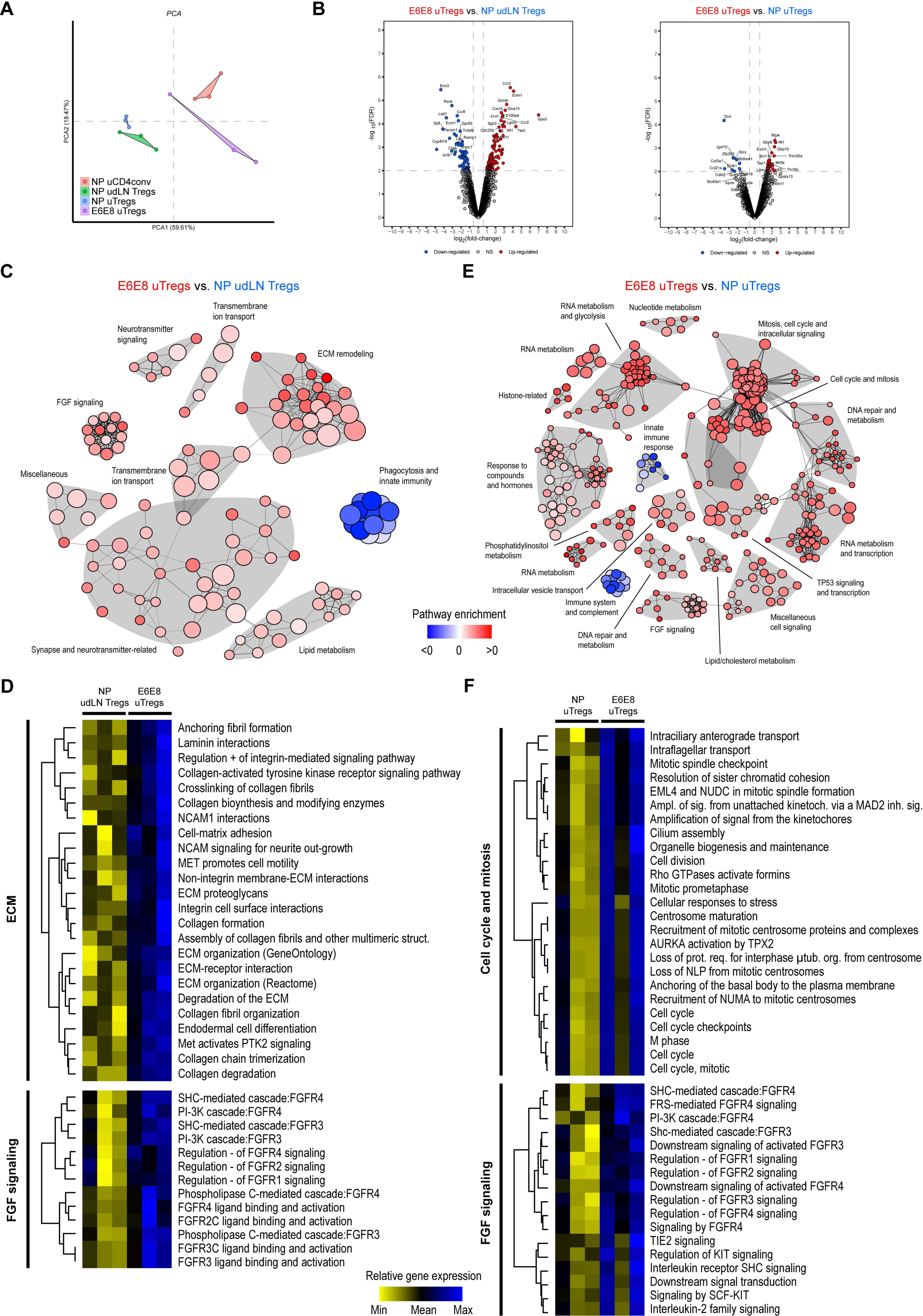
During allogeneic pregnancy the transcriptomic program of resident uTregs is tuned towards cell proliferation, trophic functions and vasculogenesis. **A)** PCA of transcriptional profiles distinguishing uTregs E-6E8, NP uTregs, NP uCD4conv and NP udLN Tregs. **B**) Volcano plot showing genes differentially expressed between uTregs E6-E8 and NP udLN Tregs (left) and between E6-E8 uTregs and NP uTregs (right). **C and E**) Network representation of pathway enrichment analysis from the list of pathways statistically differentially expressed between E6-E8 uTregs and the indicated groups. Each node in the representation corresponds to a pathway, and nodes are connected if the lists of genes in the associated pathways are overlapping. Nodes are colored using a color-gradient scale ranging from blue (<0) to red (>0) according to the difference of mean expressions between the conditions. Clusters of pathways are distinguished from each other by their grey-colored area. **D and F**) Details of the enrichment of biological pathways related to observed ECM, FGF signaling, and, cell cycle mitosis clusters in the enrichment network shown in the E6-E8 uTregs vs. NP udLN Tregs or E-6E8 uTregs vs. NP uTregs comparison. Expression score for each pathway found to be statistically differently expressed in the different individual are represented using a color-gradient scale ranging from yellow (low) to blue (high).

PCA clearly discriminated the four cell subsets (**Figure 6A**). Analysis of the top genes and pathways upregulated in E6-E8 uTregs as compared to NP udLN Tregs (**Figure 6B left**) confirmed higher expression of tissue effector Treg-related genes such as *Ccr3* and of genes associated with immune response suppression (*Ccr3, Ccr8,* and *Ccr2*), cell proliferation (*Cdc25b*, *S100a4*, and *S100a6*), blood vessel formation, development and morphogenesis (*Esm1* and *Lad1)*, and with cellular response to hormones and hypoxia (*Itga2). Cxcr3* is expressed by tissue Tregs fighting against Th1 responses (Tan et al., 2016), while *Lgmn* has been implicated in the control of *Foxp3* expression by *Garp* (Probst-Kepper et al., 2009). Likewise, compared to LN Tregs, uTregs of pregnant mice overexpress pathways related to ECM remodeling and FGF signaling (**Figure 6C** and **D)**.

There were less genes differentially expressed between uTregs from pregnant vs non-pregnant mice and they were mostly related to cell activation and metabolism (*Apoe*, *Gbp6*, *Gbp9*, *Gbp10*, and *Ifit1*), chromatin remodeling (*Bcl11b* and *Bhlhe41*), cell adhesion and migration (*Ccn1*, *Dcn*, *Col3a1*, and *Col4a1*) ), as well as angiogenesis, coagulation and uterine development (*Esm1*, *Igfbp3*, and *Igf2r*) pathways (**Figure 6B right**). Overall, the main impact of pregnancy on uTreg transcriptional profile is the overexpression of pathways related to cell-cycle, mitosis, and FGF-signaling (**Figure 6E** and **F**).

### The uTregs signature is dynamically coordinated with hypoxia and vasculogenesis during pregnancy

Many of the uTregs DEGs are associated with vasculogenesis and hypoxia. We thus investigated the link between the signatures related to Tregs, hypoxia (Moslehi et al., 2013), and vasculogenesis (Argraves and Drake, 2005) before and during pregnancy. Using a transcriptome dataset from the whole mouse uterus at different days post-conception (Nehar-Belaid et al., 2016), we applied unsupervised PCA to investigate the dynamics of these signatures in the uterine environment. We observed remarkably similar time-coordinated evolution of the uTregs (**Figure 7A right**), vasculogenesis (**Figure 7A middle**) and hypoxia (**Figure 7A left**) pathways. The two main components of the PCA, representing together more than 50% of the variance, showed a U-shaped dynamics perfectly well aligned with the duration of the pregnancy (**Figure 7A**). The statistical significance of these observations was highlighted by a very high covariance of each of these signatures and of the different stages of pregnancy (0.97). The projection of the first PCA component (PC 1) for (i) uTregs and hypoxia or (ii) uTregs and vasculogenesis showed that uTreg gene dynamics were correlated at r=0.97 with hypoxia and r=0.96 with vasculogenesis genes and followed a time-dependent progression in a well-ordered dynamic sequence from E6 to E12 (**Figure 7B**). Altogether, these results indicates that uTregs are associated with the regulation of hypoxia and vasculogenesis during pregnancy.

**Figure 7.**
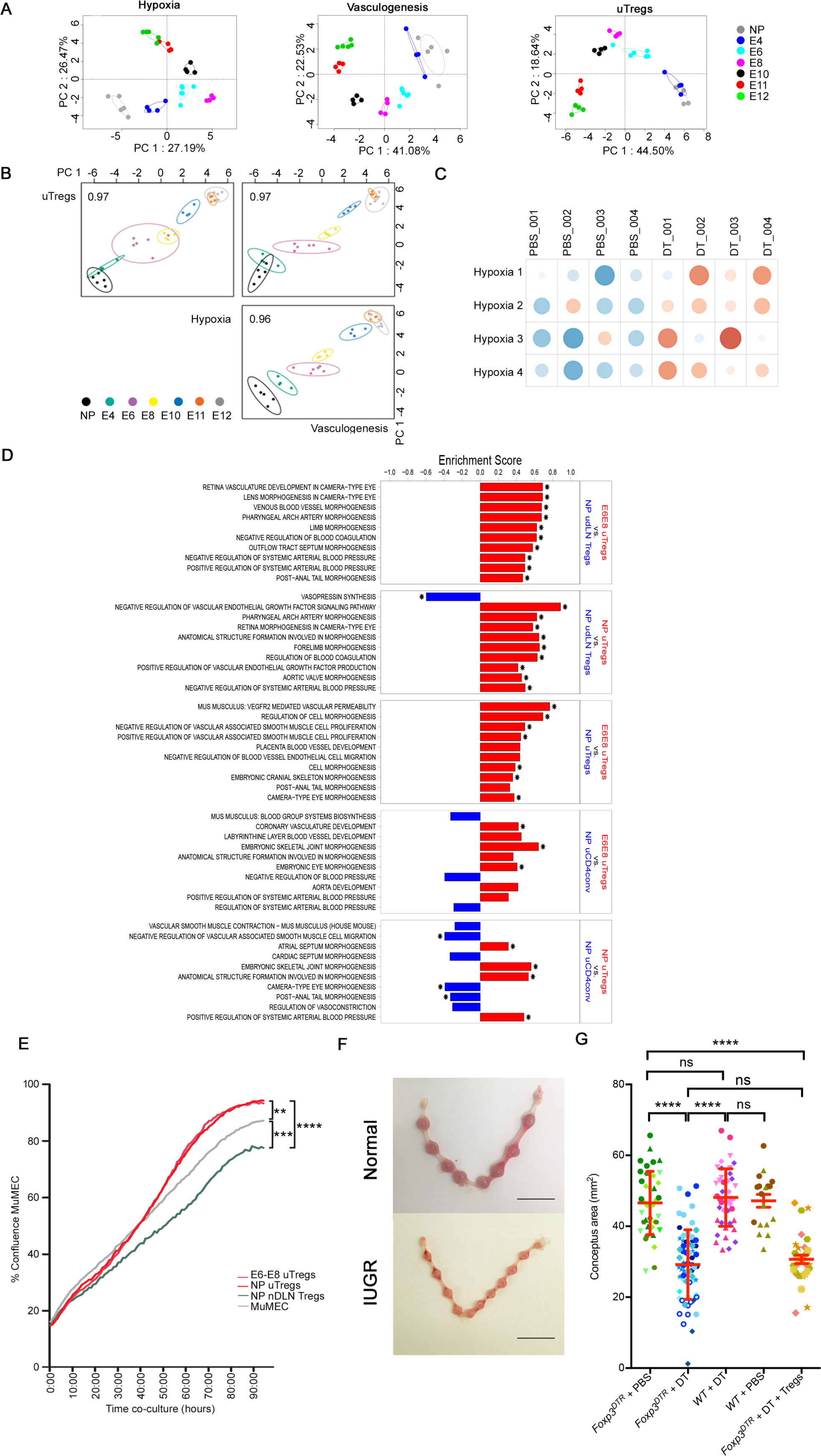
uTregs are involved in vasculogenesis, extracellular matrix remodeling and fetal survival trophic functions. **A)** Principal component analysis (PCA) based on normalized gene expression showing the coordinated dynamics of hypoxia (Moslehi et al., 2013) and vasculogenesis (Argraves and Drake, 2005) signatures with uTreg DEGs. PCA accurately separates non-pregnant (NP – E4) mice and those in early (at E6 and E8, after embryo implantation) or late allogeneic pregnancy (at E10, E11 and E12, C75BL/6 females mated with BALB/c males), (n = 4 to 6). **B)** Regularized generalized canonical correlation (RGCC) analysis for multiblock data sets representing the first component from each PCA of uTregs, vasculogenesis and hypoxia signatures from the same samples as (A). The numbers on the side are the correlation coefficients between the first components extracted from each signature. The colors and ellipses are related to the pregnancy time points and indicate the power of each component to discriminate between the different stages of pregnancy. **C)** Differential transcriptional analysis of total uteri after Treg depletion in the Foxp3^DTR^ mouse line. Transcriptomic module analysis of Treg sufficient or Treg depleted total uteri (n = 4 per group, see the Methods section for the depletion protocol). Modules shows gene clusters from molecular signatures related to blood vessel formation and development, and extracellular matrix functions. Cluster labels represent MSigDB molecular signatures from C2 (curated gene sets) and C5 (GO gene sets). The average expression over all genes in each module is shown per sample (rows) colored from red (over-expressed) to blue (under-expressed). Module expression correlates with sample status (DT vs. PBS) (chi-square test p-value < 5e^-2^) and is significantly different between the groups (t-test p-value < 1e^-2^). **D)** Enrichment scores of vascular system-associated pathways identified in the different lists of differentially expressed genes. Significant items are represented with a star and are colored based on the directionality of the dysregulation. **E)** Confluence assay during 90h co-culture between uTregs (red), uDLN (blue) or non-DLN (green); and with mouse primary uterine microvascular endothelial cells (muMECs). Data represent 2 different experiments, 4 points, 10 E6-E8 pregnant BALB/c-mated or NP C57BL/6 females per point. Significant differences are denoted by **: P < 0.01, ***: P < 0.001 and ****: P < 0.0001. **F)** Conceptus area micrography of normal gravid uterus at E8 or gravid uterus with fetuses presenting intrauterine growth restriction (IUGR), respectively from PBS or DT-treated FoxP3*^DTR^* mice (C57BL/6 background mated with BALB/c males) (scale bar = 1cm, n=5). **G)** Scatter dot plot of conceptus area at E8 from WT (C57BL/6) or Foxp3*^DTR^* mice that received PBS or DT every 2 days from E1,5, or DT every 2 days from E1,5 plus 5-9 × 10^6^ T reg cells from WT mice. Each littermate is distinguishable by a distinct color and shape (Foxp3^DTR^ + PBS: n = 6 littermates; Foxp3^DTR^ + DT: n= 8; WT + DT: n = 5; WT + PBS: n = 2; Foxp3^DTR^ + DT + Tregs: n = 5). Data represent 198 embryos in total from C57BL/6 or FoxP3*^DTR^* mice mated with BALB/c males (7 experiments). Red lines represent mean and standard deviation. A Student’s t test was used for statistical analysis. ns: non-significant; ****: P < 0.0001.

### uTregs act as positive regulators of vasculogenesis favoring proper fetal-placental growth

To test the hypothesis that uTregs act as positive regulators of vasculogenesis favoring proper fetal growth, we first investigated the changes in uterine vasculogenesis signatures in the presence or absence of Tregs. Using the Foxp3^DTR^ mouse model (**Figure S5A**), we generated a complete transcriptome from uterine tissues at E8 from control-(PBS) and diphtheria toxin-(DT) treated mice (**Figure S5B**). Compared to DT-treated mice, in the presence of Tregs, we found a positive enrichment for GO terms related to embryonic and blood vessel development (**Figure S5C**). Furthermore, using functional modules assembled from more than 100 molecular signatures of hypoxia, vasculogenesis, angiogenesis and extracellular matrix remodeling functions from the Molecular Signatures Database (MSigDB) (**Figure S5D**), we observed that the four top modules that discriminate the Treg-depleted uterus are related to “hypoxia”, and are all up-modulated in the absence of Tregs (**Figure 7C**). This is consistent with the elevated expression of biological pathways related to the vascular system and angiogenesis in uTregs from pregnant or not pregnant mice as compared to uterine helper T cells or Tregs from PALN (**Figure 7D**). Collectively, these results further support that uterine Tregs are involved in the process of vasculogenesis and prevent hypoxia of the uterus.

These observations led us to evaluate the direct trophic effects of uTregs. We first investigated the direct effects of uTregs on placental-derived endothelial cell growth. Mouse primary uterine microvascular endothelial cells (muMECs) were grown in the presence of LN Tregs from NP mice or uTregs from NP or E6-E8 mice, and were time-lapse imaged over 90 hours. uTregs significantly improved the proliferation of muMECs independent of the pregnancy status of the mice. In contrast, LN Tregs had a negative effect on muMECs’ proliferation (**Figure 7E**).

As impaired placental angiogenesis leads to intra-uterine growth retardation (IUGR) (Ahmed and Perkins, 2000), we measured conceptus number and size in the presence or absence of Tregs. We investigated the effects of Treg depletion on the conceptus at E8 because, with our mice and in our animal facility, the scheme of DT administration and its timing did not result in severe systemic inflammation at E8, as described in (Kim et al., 2007). Treg depletion after mating did not affect implantation numbers at E8 and there was no ongoing inflammation of the conceptus, indicating that fetal immune rejection had not yet occurred. Actually, the Treg-depleted pregnant females had no obvious signs of a systemic disease at the time we analyzed them, despite major Treg depletion. However, in the absence of Tregs, we observed a highly significant and marked IUGR (**Figures 7F-G**).

To ascertain that IUGR was caused by depletion of uTregs, and not circulating Tregs, we sought to selectively ablate resident uTregs. Taking advantage of the slow repopulation of uterus by donor Tregs (see **Figure 1**), we adoptively transferred 5 to 9 × 10^6^ wild type Tregs from C57BL/6 mice at the time of the first DT injection. This replenished the spleen and the para-aortic lymph nodes with Tregs, but not the resident uTregs compartment (**Figure S5E**), generating mice with a specific uTregs depletion. A very significant IUGR was still observed in these mice (**Figure 7F**). This indicates that resident uTregs are specifically required for proper fetal-placental growth, at least until E8 of pregnancy.

## DISCUSSION

### A novel population of resident effector memory Tregs in uterine tissue

We have identified a novel Treg population with an effector memory phenotype that resides and self-maintains in the uterine tissue. This population can be defined as the *bona fide* uterine Tregs based on blood wash-out experiments that discriminate intra-tissue Tregs from tissue-blood Tregs. Resident uTregs from the non-pregnant uterus represent approximately 10% of the uterine CD4^+^ T cells. The uTreg phenotype is comparable to that of resident effector memory T cells (Trm) in non-lymphoid tissues, with the expression of CD69, CD103, IL33R. uTregs do not express the chemokine receptor CCR7 (Smigiel et al., 2014) and show slow recirculation in parabionts. uTreg expression of Ki67 and EdU incorporation profile both indicate extensive proliferation. Comparison of the transcriptomes of uTregs and uterine CD4^+^ Tconv highlights a clear Treg signature. Consistently, uTregs are almost as suppressive as splenic Tregs in a miniaturized suppression assay. Thus, there is a previously unidentified population of Tregs that resides and self-maintains within the uterus.

### uTreg homeostasis

In non-pregnant mice, uTregs localize in the endometrium and gather around the glandular uterine epithelial cells. Upon embryo implantation, uTregs upregulate their expression of CCR10 and CXCR5, the chemokine receptors for CCL28 and CXCL13 respectively. CCL28 is expressed by endometrial cells (Choi et al., 2016), while CXCL13 levels are increased in the plasma and decidua during pregnancy (Nhan-Chang et al., 2008) and expressed by decidual cells (Munoz-Fernandez et al., 2012) and in the endometrium (Franasiak et al., 2015). Thus, uTregs have an enhanced expression of molecules that could drive uterine tissue localization. Remarkably, uTregs survive and proliferate in an environment deprived of IL2 and are thus IL2-independent contrary to Tregs from lymphoid organs. The fact that they rapidly re-acquire CD25 expression upon IL2 administration indicate that the residency in the uterus is the cause for loss of CD25 expression. TGF-β, IL-15 and IL-33 are likely collectively supporting uTreg homeostasis as (i) they can each support Treg survival (Vang et al., 2008), (Kuswanto et al., 2016), (ii) the uterine milieu contains high levels of these cytokines, and (iii) uTregs express their receptors. While our transcriptomic analysis does not show an upregulation of the LH, CG or progesterone receptors in uTregs, we cannot exclude that these cells express these receptors, which could then be implicated in recruiting Tregs during pregnancy (Schumacher et al., 2013, 2014, 2017; Areia et al., 2015).

### Comparison of uTregs with other tissue-resident ST2^+^ Tregs

A large fraction of tissue-resident Tregs has been described as ST2^+^ (IL33 receptor) in infected lung, injured muscle, visceral adipose tissue (VAT) and colon (Arpaia et al., 2015; Burzyn et al., 2013b; Feuerer et al., 2009; Panduro et al., 2016; Schiering et al., 2014). ST2^+^ tissue-Tregs are phenotypically described as CD44^high^CD62L^low^PD-1^high^, with some also expressing KLRG1 and CD103 (Arpaia et al., 2015; Schiering et al., 2014; Vasanthakumar et al., 2015), and ST2^+^ Tregs from muscle or lung express amphiregulin (Burzyn et al., 2013b; Feuerer et al., 2009). uTregs also express ST2, amphiregulin and effector memory markers and thus have a phenotype reminiscent of ST2^+^ tissue-specific Tregs. However, the uTreg transcriptional profile is distinct from those of colon, VAT and muscle tissue Tregs, notably for the uTregs DEGs involved in trophic function.

### Antigen specificity of uTregs

The TCR repertoire of uTregs from non-pregnant mice is polyclonal. This argues against a residency that would be triggered mainly by tissue-specific antigens (Li et al., 2021). Upon pregnancy, uTregs highly proliferate. This leads to a TCR repertoire that is still polyclonal, but with a few clonotypes that have major expansions. While the sharing of NP uTreg TCRs with TCRs from SLOs is around 1%, the sharing of uTregs TCRs from pregnant mice with these raises to 3-4%. In addition, these shared TCRs have a high generation probability (Pgen). This indicates that the expanded uTregs from pregnant mice have more public-TCRs. We thus hypothesize that uTregs residency is more likely triggered by the uterine tissue-specific microenvironment that would trigger bystander cells to initiate residence. These cells are then expanded during pregnancy, with a preferential expansion of the cells expressing public TCRs. Further single-cell studies should help refine these conclusions.

### The singular trophic and tissue-remodeling phenotype and function of uTregs

Transcriptome studies reveal that, already before pregnancy, uTregs upregulate several genes involved in uterine tissue maintenance, extracellular matrix (ECM) remodeling and blood vessel development. This suggests a role of Tregs in the remodeling of the uterus during the estrus/menstrual cycle. Upon pregnancy, uTregs maintain this remarkable transcriptome profile, which is not shared with other tissue-specific Tregs.

Furthermore, our unsupervised analyses of the uTregs, hypoxia and vasculogenesis signatures of the uterine environment revealed a remarkable coordination of the dynamics of uTregs with hypoxia or vasculogenesis signature dynamics. PCA distinguished three different transitions during pregnancy: non-pregnant and pre-implantation E4, peri-implantation E6-E10 and middle pregnancy E11-E12. We explored this link between the uTregs and vasculogenesis or hypoxia transcriptome signatures during pregnancy with genes and pathways-based statistical methods. We found a strong correlation between the first components of the regression model between uTregs gene expression dynamics and that of genes involved in vasculogenesis and hypoxia. This is another indication of the role of uTregs in vasculogenesis during pregnancy.

### uTregs modulate trophic function and conceptus growth

The role of uTregs in vasculogenesis during pregnancy was further confirmed by the effect of Treg depletion on the uterine microenvironment at E6-E8. Compared to the unmanipulated uterine transcriptome, the transcriptome of uteri from Treg-depleted mice revealed upregulation of genes in response to hypoxia and downregulation of genes involved in embryonic and blood vessel development. These two functions appeared to be specifically enhanced in uTregs compared to other Tregs (**Figures 6 and 7**). The indirect transcriptomic evidence for uTregs involvement in blood vessel development were strongly supported by *in vitro* cell proliferation assays showing that uTregs, but not other Tregs, enhance microvascular primary endothelial cells proliferation (**Figure 7)**.

Finally, we determined in a model of uTreg deficiency that the size of the conceptuses from uTreg-depleted mice was markedly and significantly decreased at E6-E8, confirming a direct and specific role of uTregs in embryonic development. These observations are supported by a recent study wherein Treg depletion at E7 resulted in increased uterine artery resistance caused by vasoconstriction (Care Alison S. et al., 2018). Thus, our and other recent studies indicate that uTregs have an essential role in modulating uterine artery function during early and middle pregnancy. These results are supported by the associations between abnormal lymphatic vessel development and decreased decidual regulatory T cells in patients with severe preeclampsia (Jung et al., 2018).

Based on these findings, we propose a role of Tregs in pregnancy that can be separated into 3 phases. During the first phase, rapid response of uTregs in combination with a systemic response of self-specific memory Tregs in udLNs prevents the initiation of an immune response that could be triggered by release of cell debris due to high proliferation of endometrial tissue. During a second phase, uTregs contribute to the increased need for uterine tissue remodeling for pregnancy and placentation; the modest numbers of uTregs suggests that they could orchestrate the function of other immune and non-immune cells such as NK cells, which are more prevalent than uTregs. In a third phase, accompanying fetal mass increase, pTregs could be enlisted to help control the anti-fetal immune response (Samstein et al., 2012). Altogether, our work further highlights the “*chameleonesque”* and versatile functional properties of Tregs and suggest new therapeutical schemes to prevent the risks of IUGR in preeclamptic patients.

## ACKNOWLEDGMENTS

We thank Guillaume Churlaud, Felipe Leal-Valentim, Federica Martina, Oceane Konza, Xinyue Li and Ariadna Gonzalez-Tort who had various input in this work. We thank Christelle Enond, Flora Issert, Kim Nguyen, Olivier Bregerie, and Bocar Kane (Centre d’Exploration Fonctionnelle, Université Pierre et Marie Curie) for animal care, Bruno Gouritin for cell sorting by flow cytometry, Tudor Manoliu for Imaging and Vanessa Mhanna for a TCR dataset. This work was supported by a grant from the European Research Council Advanced Grant (ERC⍰2012⍰AdG, TRiPoD, Agreement number 322856, to D. Klatzmann), and LabEx Transplantex, Strasbourg, France (to N. Mooney). LM. Florez was supported by a DIM Biotherapies ILE DE FRANCE Region grant.

## AUTHOR CONTRIBUTIONS

DK conceived, designed, and supervised the study and obtained funding. LMF, GF, GDJ, DNB, TC, MGR, and JL acquired the data. LMF, GF, GDJ, PB, KGF, PR, SZ, MG, JL, NM, NT, EMF and DK analyzed and interpreted the data. LMF, GDJ, PB, PR, NT performed the statistical analyses. LMF, GF, SZ and MG provided technical and material support. LMF, GDJ, NT, EMF and DK drafted the manuscript. All authors critically revised the manuscript for important intellectual content.

## DATA SUBMISSION

Murine datasets of purified Tregs and the CD4conv cell RNA-Seq transcriptome were deposited in the GEO repository (GSE109895).

Murine datasets of the total uterine microenvironment transcriptome after Tregs depletion in the Foxp3^DTR^ model were deposited in the GEO repository (GSE123463).

## METHODS

### Mouse strains

C57BL/6 (B6) and BALB/c female mice were from Janvier SAS. *Foxp3EGFP (Foxp3^eGFP^)* (Wang et al., 2008), Foxp3-GFP-DTR ‘knock-in’ (Foxp3^DTR^) (Kim et al., 2007) and RAG^-/-^ all from a B6 background were previously described. BALB/c background Foxp3^IRES-GFP^ and Foxp3^DTR^ females were used for the uTreg and total uterus Treg depletion RNA-seq analysis, respectively. Animals were maintained in our animal facility under specific pathogen-free conditions in agreement with current European legislation on animal care, housing, and scientific experimentation. All procedures were approved by the Regional Ethics Committee on Animal Experimentation No. 5 of the Ile-de-France region (Ce5/2012/031).

### Timed mating

Estrous phases of adult females were determined by the vaginal cytology method. One or two eight-week-old female mice in estrus were set up in the afternoon with individual males for mating. All matings performed in the study were allogeneic matings between BALB/cJ males and females from the C57BL/6J background, except if stated otherwise. Females were checked daily for the presence of a vaginal plug in the morning; the day of plug detection was considered as day E1. Plugged females were analyzed at days 0, 6, 7, 8, 9, 10, 11 and 12 after mating. Most non-pregnant females were in estrus. Naïve and pregnant females were euthanized by cervical dislocation.

### Immunofluorescence

*Foxp3^eGFP^* non-pregnant and pregnant females at E8 were deeply anesthetized with ketamine/xylazine and transcardially perfused with PBS,, followed by 10⍰mL of 4% paraformaldehyde (PFA) (Electron Microscopy Science #15710). Uteri, draining para-aortic lymph nodes and non-draining brachial lymph nodes were harvested and post-fixed overnight in 4% PFA. Fixed organs were embedded in 30% sucrose for 48 h and dissected into 80 μm thick sections with a freezing microtome. Transversal and longitudinal sections were collected. Tissue sections were blocked for 30 min with 0.1 M lysine, and stained overnight with primary antibody cocktails diluted in the blocking-permeabilization solution 0.1 M lysine - 1% Triton (Tx) – 0.1% pork gelatin – FBS 3% PBS 1X. Samples were further stained for 1 h with the secondary antibodies. Sections were mounted with Prolong Gold anti-fade Reagent (Invitrogen, P36930) on microscope slides (Superfrost plus, Thermo Scientific J18000AMNZ) and covered with coverslips (Menzel Gläser, Thermo Scientific). After 48 h, the edges of the slides were sealed with nail polish. The primary antibodies used were: polyclonal chicken anti-mouse GFP (abcam 13970, dilution 1:1000) and rabbit monoclonal anti-FOXA2 (abcam 108422, dilution 1:200). The secondary antibodies used were: Alexa-Fluor 488 conjugated goat anti-chicken polyclonal (abcam 150173, 1:1000) and Alexa-Fluor 633 conjugated goat anti-rabbit (Life Technologies A21072, 1:1000). Images were acquired with a Leica SP5 confocal microscope and processed with Fiji ImageJ software (Plateforme d’Imagerie ICM Hôpital Pitié Salpêtrière).

### Flow cytometry

Single-cell suspensions from lymph nodes and spleen were prepared by mechanical disruption after dissection. Uteri were isolated and carefully dissected to remove the yolk sac containing the fetus followed by mechanical disruption and digestion with DNase I 200 units/mL (Roche 10104159001) and Liberase DL 0.3 W units/mL (Roche 05466202001) in RPMI (Gibco 31870025) for 25 min at 37°C.

Samples were blocked using anti-CD16/CD32 (supernatant of hybridoma 2.4G2; American Type Culture Collection, 1:40), followed by the addition of fixable viability dye eFluor 780 (LD) (eBioscience 65056518, 1:1000) for 20 min at 4°C. Fluorophore-conjugated antibodies were from BD Biosciences and eBioscience: R-phycoerythrin Texas red-labeled (PETR) anti-CD45 (1:1600); efluor 450 (ef450) or R-phycoerythrin-labeled (PE) or violet Horizon-labeled (v500) anti-CD3 (1:400); v500 or allophycocyanin-labeled (APC) anti-CD4; AlexaFluor700-labeled (A700) anti-CD8 (1:400); PE-cyanin 7-labeled (PE-Cy7) or A700-anti-CD25(1:200); PE-anti-CTLA-4 (1:200); PE-or APC-anti-IL15Rα; PE- or APC-anti-GARP; PE-anti-IL33R; FITC- or PeCy7-anti-CD44 (1:200); ef450- or ef780-anti CD62L (1:200); PERCP-anti-CD90.1 (1:200); APC-anti CD90.2 (1:200). PeCy7-anti-CCR7 (1:50) staining was performed for 30 min at 37°C prior to cell surface staining.

Uterine staining of resident cells was performed in anesthetized (ketamine/xylazine) non-pregnant females transcardially perfused with 50 mL of PBS1X. Intracellular staining was performed using the FOXP3 mouse Treg cell staining kit (eBiosciences 00552300). Intracellular labeling of FOXP33, Ki67 and CTLA-4 was done with a PE-, APC-, eF450- or FITC-labeled anti-mouse FOXP3 antibody (FJK-16s; eBioscience); FITC- or PE-anti Ki67 (1:1000); PE-anti-CTLA-4 (1:200). Stained cells were analyzed using an LSRII flow cytometer (BD Biosciences), and data were analyzed using FlowJo software (BD Biosciences).

### Uterus quantitative RT-PCR

Uterine segments were carefully dissected to remove the yolk sac containing the fetus, immersed in 0.5 mL of RNA-later (Qiagen, Hilden, Germany) and stored at -80°C. For RNA extraction, total tissues were transferred into 1 mL of TRIzol Reagent (Life Technologies, France), and disrupted and homogenized with a TissueLyser II (Qiagen). Total RNA was isolated by using RNeasy Mini Kits (Qiagen). cDNA was generated using SuperScript III (Life Technologies) according to the manufacturer’s instructions. Quantitative PCR was performed using the 7500 Fast Real-Time PCR System (Life Technologies) with Fast Master mix (Life Technologies). The probe IDs were: TNF-a: Mm00443258_m1, IFN-g: Mm01168134_m1, IL4: Mm00445259_m1, IL10: Mm00439614_m1, IL15: Mm00434210_m1, IL33: Mm00505403_m1, TGF-β: Mm01178820_m1, 18S: Mm03928990_g1, and GAPDH: Mm99999915_g1. PCR was performed in triplicate and the mRNA levels were normalized to that of GAPDH and to 18S mRNA levels. Gene expression in non-pregnant and E8 or E12 pregnant groups was expressed as relative transcript abundance and fold increase relative to the control group.

### ELISA assay

To measure murine IL2 levels, we harvested total uteri, udLN and ndLN samples from non-pregnant females. Samples were mechanically disrupted. Uteri were digested as previously described. Cells were plated in 96-well plates (Nunc, Denmark) at 5×10^5^ cells/well and stimulated with phorbol 12-myristate 13-acetate (PMA) (Sigma P8139; 50 ng/mL) and ionomycin (Sigma I0634; 500 ng/mL) in RPMI for 4 h at 37°C and 5% CO_2_. Supernatants were collected and frozen and kept at -80°C until use. Secretion level of IL2 was measured according to the manufacturer’s recommendations using the specific mouse IL2 ELISA Ready-SET-Go kit (eBioscience 88-7024-88). The OD was read at 450 nm using an automatic ELISA plate reader (DTX 880, Multimode detector; Beckman Coulter).

### *In vivo* Tregs depletion

Foxp3^DTR^ female mice (C57Bl/6 background) were mated with BALB/c males and treated every second day with intraperitoneal injections of 1 µg of diphtheria toxin or PBS1X at E1, E3, E5, and E7 days of pregnancy. That treatment regimen allowed the depletion of 85% of total Tregs after four injections and was based on the study done for the Foxp3^DTR^ mouse line (Kim et al., 2007).

### AAV-IL2 treatment

Six-week-old C57BL/6 mice were injected intraperitoneally with 10^12^ viral genomes (vg) of recombinant AAV8 vectors carrying the transgenes for luciferase (AAV-LUC) and murine IL2 (AAV-IL2) as previously described (Jimenez et al., 2011; Churlaud et al., 2014). After 14 days, non-pregnant females PBS-perfused uterus, udLNs and ndLNs were harvested for flow cytometry analysis.

### EdU proliferation assay

To measure the proliferation ability of uTregs, we used the Click-iT® Plus EdU Flow Cytometry Assay Kit (Life Technologies) and followed the manufacturer’s protocol. EdU (5-ethynyl-2’-deoxyuridine) is a nucleoside analog to thymidine and is incorporated into DNA during active DNA synthesis. Detection was based on a click reaction, a copper catalyzed covalent reaction between a picolyl azide and an alkyne. The picolyl azide was coupled to Alexa Fluor® 647 dye. The percentages of S-phase Treg cells in the uterine, udLN and ndLN populations of non-pregnant C57BL/6 females were determined by flow cytometry.

### Parabiosis

Female 6- to 8-week-old congenic Foxp3^IRES-GFP^ BALB/cJ CD90.1 (GFP+ CD90.1) and BALB/cJ CD90.2 (GFP-CD90.2) mice were surgically connected in parabiosis in couples GFP^+^/GFP^-^ as previously described (Kamran et al., 2013). After corresponding lateral skin incisions were made from elbow to knee in each mouse, forelimbs and hindlimbs were tied together using nylon suture, and the skin incisions were closed using stainless steel wound clips. After surgery, mice were given buprenorphine every 12 h and maintained on a diet supplemented by Bactryl for prophylaxis of infection and distress. 14 days after surgery, flow cytometry analysis was performed in each parabiont. The percentages of CD90.1 and CD90.2 among CD8^+^, CD4^+^Foxp3^-^ and CD4^+^Foxp3^+^ T cells in the uterus, udLNs, parabiosis dLNs (pbdLNs), non-dLNs (ndLNs), and spleen were used to calculate the host-to-donor ratio in each parabiont.

### Treg cell sorting of peripheral CD4 and Tregs

Non-pregnant Foxp3^eGFP^ and C57BL/6 females were used to isolate peripheral CD4^+^ or Treg cells by cell sorting using FACS ARIA (BD Biosciences). Total lymph nodes and spleens were dilacerated, and cells were pooled from 5 to 10 mice. A negative enrichment in CD4 cells was performed using a biotinylated antibody cocktail: Ter-119 (1:400), CD11c (1:200), CD11b (1:200), CD8 (1:200) and B220 (1:400) (all from BD Biosciences) and anti-biotin magnetic micro-beads (Miltenyi Biotec 130095485). The solutions were sorted with the AutoMACS (Mitenyi Biotec) and the negative fraction was immunostained for FACS ARIA cell sorting: PETR-anti-CD45 (1:1600), APC-anti-CD4 (1:200), PE-Cy7- or Af700–anti-CD25 (1:200), PeCy7-anti-CD44 (1:200) and ef450-anti-CD62L (1:200). Cells were used for adoptive transfer experiments.

### Adoptive cell transfers

FACS ARIA purified CD4+Foxp3-GFP-CD44-CD62L+ T cells from NP Foxp3^eGFP^females were injected i.v. into C57BL/6 or RAG^-/-^ non-pregnant females at 6×10^6^ cells per mouse. 2 days after injection the females were progressively mated to BALB/c males and the uteri were analyzed after 15 days on days 14, 13, 12, 10, 8, 6 and 0 of pregnancy. Mesenteric lymph nodes were used as controls of Nv CD4 T cell injection and induction of Foxp3.

94% pure CD4^+^CD25^hi^ Tregs were isolated via triple column selection on an autoMACS® Pro Separator (Miltenyii Biotec) by using the *Mouse CD4^+^ CD25^+^ Regulatory T Cell Isolation Kit* (Miltenyii Biotec) according to the manufacturer instruction. Then 5 to 9 × 10^6^ cells were injected i.v. into plugged Foxp3*^DTR^* mice at E1.5.

### Proliferation / Suppression Assays

CD4^+^Foxp3^+^ Tregs, CD4^+^Foxp3^-^ helper T cells and CD11c^hi^ dendritic cells were sorted from the spleen and uterus of naive Foxp3^IRES-GFP^ mice. Helper T cells were labelled with VPD-450 (Invitrogen) according to the manufacturer instructions. Then, 2000 Helper T cells were cultured at 37°c in 96-round bottom well plates in the presence of 600 dendritic cells and 1 μg/mL of anti-CD3ε antibody ± 500 Tregs sorted from the spleen or uterus of naive Foxp3^IRES-GFP^ mice. VPD_450_ dilution of helper T cells was measured by flow cytometry after 3 days.

### Endothelial cell proliferation assay

C57BL/6 mouse primary uterine microvascular endothelial cells or muMECs were from Cell Biologics (distributed by Euromedex, Souffelweyersheim, France). These cells were cultured at 37°C, 5% CO2 and 95% air in a humidified incubator, between passages 3 and 7 in complete mouse endothelial cell medium (from Cell Biologics, Euromedex) composed of mouse endothelial cell medium with the addition of endothelial cell medium supplement kit (from Cell Biologics, Euromedex) containing fetal bovine serum, antibiotics, glutamine, hydrocortisone, epidermal growth factor or EGF, heparin, endothelial cell growth supplement or ECGS and vascular endothelial growth factor or VEGF. The muMECs were seeded in T75 tissue culture flasks pre-coated with gelatin-based coating solution for 2 min (from Cell Biologics, Euromedex) and cultures were divided twice a week or as necessary. Non-pregnant and pregnant E6-E8 BALB/c background Foxp3^IRES-GFP^ females were used to isolate uterine, udLN and non-draining (ndLN) lymph nodes by cell sorting using FACS ARIA (BD Biosciences). Groups of 10 mice per NP sample and 8 mice per pregnant sample (E6-E8) were made and 4 samples were collected per cell type. Total uteri and draining lymph nodes were digested with DNase I 200 units/mL (Roche 10104159001) and Liberase DL 0.3 Wunits/mL (Roche 05466202001) in RPMI (Gibco 31870025) for 25 min at 37°C. The solution was immunostained for FACS ARIA cell sorting: PETR-anti-CD45 (1:1600), APC-anti-CD4 (1:200) and PE-Cy7-anti-CD25 (1:200). A positive enrichment in CD4^+^ cells was performed using the EasySep Mouse APC Positive Selection Kit (Stemcell 18452) and Stemcell^TM^ Magnet.

Coculture experiments were carried out with muMECs (1000 or 1500 cells/well) and purified Tregs at a ratio 1:1 in 96-well plates pre-coated with gelatin-based coating solution (from Cell Biologics, Euromedex).

Co-cultures were immediately placed in a humidified incubator and real-time monitoring was commenced. Confluence of the muMECs was monitored in real-time using Incucyte® technology (Essenbio science, Ann Arbor, MI, USA). Images were acquired every hour for 90 h and real-time evaluation of muMEC confluence was analyzed by Incucyte® S3 software.

#### RNA-Seq profiling of Tregs or Tconv cell populations extracted from uterine tissue or uterus-draining lymph nodes of pregnant or non-pregnant mice

*Groups*: non-pregnant and pregnant E6-E8 BALB/c background Foxp3^IRES-GFP^ females were used to isolate by FACS (BD FACSAria) several cell populations: uterine Tregs from non-pregnant mice (NP uTregs), udLN Tregs from non-pregnant mice (NP udLN Tregs), uterine CD4conv from non-pregnant mice (NP uCD4conv) and uterine Tregs from pregnant mice (E6-E8 uTregs).

##### Samples preparation

to ensure a sufficient yield of target cell populations, each sequenced sample (n = 3 per group) was generated using a pool of mice (20 or 3-8 for cell populations coming from non-pregnant and pregnant mice, respectively). Total uteri and draining lymph nodes from mice perfused with PBS before organ collection were digested with DNAseI 200 units/mL (Roche 10104159001) and Liberase DL 0.3 Wunits/mL (Roche 05466202001) in RPMI (Gibco 31870025) for 25 min at 37°C. The solution was immunostained for FACS with PETR-anti-CD45 (1:1600), APC-anti-CD4 (1:200) and PE-Cy7-anti-CD25 (1:200). A positive enrichment in CD4+ cells was performed using the EasySep Mouse APC Positive Selection Kit (Stemcell 18452) and StemcellTM Magnet. RNA was purified with the RNAqueous®-Micro Total RNA Isolation Kit (Invitrogen AM1931). cDNA was prepared with the SMART-Seq v4 Ultra Low Input RNA Kit. Indexed paired-end libraries were done with the Nextera® XT Library Prep Kit and the Nextera XT Index Kit. The quality control was done in Tapestation and Quantifluor. Illumina sequencing of the 12 samples in 2×150, 50 million reads/sample was done with the NextSeq 500 High Output Kit v2 (300 cycles) sequencer (Illumina Inc) generating paired end reads of approximately 2 × 150 bp in length. Murine datasets were deposited in the GEO repository (GSE109895).

##### Data transformation and normalization

Gene counts were transformed and normalized using *DESeq2* R package and VST method. Then, batch effects were removed with *ComBat* method using “tissue” and “sample group” of origin as covariates.

##### Gene sets overexpression analysis

we performed pathways overexpression analysis using pathways from Gene Ontology (GO), KEGG, PANTHER, Reactome and WikiPathways databases with *GSVA* R package. Then, the differential analyses between the groups of interest were performed using standard *limma* R package, following authors instructions.

##### Networks of gene sets

we computed pairwise the similarity between the significantly overexpressed pathways (p-value ≤ 0.05). The similarity is defined here as the percentage of common genes between two gene sets. We used this similarity matrix to create the network graphs using *igraph* R package. On a network, each node represents a gene set, which are connected if the similarity value between two of them is greater than a defined threshold (typically within the range 0.1-0.2). Next, we removed the nodes which were not connected to any other one as well as clusters containing too few pathways (usually between 3-5). Finally, we determined clusters of pathways around highly connected and dense node clouds and extracted their contents in gene sets. We finally used this information to manually annotate every cluster for each network.

#### RNA-Seq profiling of uterine tissue from pregnant Foxp3^DTR^ mice treated with DT or PBS

##### Samples preparation

allogenic Foxp3^DTR^ females were mated with BALB/c males and treated with DT as described above. Females were sacrificed at day E8 of pregnancy and the entire uterus (with fetal tissues) was collected, incubated overnight in RNAlater (Qiagen) at 4°C and then transferred to -80°C for storage. Samples were then lysed and homogenized using the tissue lyser (Qiagen) and total RNA was purified using Trizol (Invitrogen) according to the manufacturer’s instructions. RNA yield was assessed using a NanoDrop 1000 spectrophotometer (NanoDrop Products, Thermo Fisher Scientific). RNA integrity was assessed using an Agilent Bioanalyzer showing a quality RNA integrity number of 8-10 (Agilent Technologies). The RNA was processed using the Illumina TotalPrep RNA Amplification Kit Protocol according to the manufacturer’s protocol. High-quality libraries were prepared using the Kapa mRNA hyperprep kit (Roche). The quality control was done in Tapestation and Quantifluor. Illumina sequencing of the 8 samples in 100 million reads/sample was done with the NextSeq 500 High Output Kit v2 (300 cycles) sequencer (Illumina Inc) generating paired end reads of approximately 2 × 75 bp in length. Datasets were deposited in the GEO repository (GSE123463).

##### GeneOntology pathways enrichment analysis

using this dataset, we also performed GeneOntology (GO) pathway enrichment analysis based on differentially expressed genes in between the two groups using the R package clusterProfiler (Yu, Wang, Han, & He, 2012). From the GO “biological process” branch, we selected the root node and the sub-nodes “immune system process” (GO:0002376), “developmental process” (GO:0032502), “regulation of reproductive process” (GO:2000241) and “system development” (GO:0048731), as well as their respective “children” and “grand-children” nodes. We considered as enriched the terms showing a p-value < 1e^-2^ and the ones that appear among the top five on the list ranked in each GO tree category.

#### Quality control and handling of RNA-Seq raw data

For all our RNA-Seq datasets described earlier, we first generated quality control reports of the raw fastq files using FastQC. Then, MultiQC (Ewels et al., 2016) summaries over all samples were created, which reported show satisfactory quality control. The Nextera adaptor sequences were removed using Trimmomatic (Bolger et al., 2014). The trimmed reads of each sample were aligned against the Mus musculus genome using default parameters of Salmon for pair ended sequences. The genome index database was built using the Ensembl genome (Yates et al., 2016) version GRCm38. Next, featureCounts (Liao et al., 2014) was used with default parameters for forward stranded sequences to count the reads that align with genomic features of interest. The annotation file used here was also downloaded from the Ensembl database (version Mus_musculus.GRCm38.87.chr.gtf). The analysis was performed at gene level. The read counts table was further analysed with R scripts following a workflow designed specifically for this dataset. We proceeded with the unambiguous read counts at gene level.

#### Microarray profiling of uterine tissue from non-pregnant mice or at different pregnancy stages

##### Samples preparation

uteri from non-pregnant mice (control samples): n=6 and allo-pregnant E4: n=4, E6: n=6, E8: n=4, E10: n=4, E11: n=4 and E12: n=5 C57BL/6 mice were used to generate the total uterine microenvironment transcriptome. Murine datasets were published in the GEO repository (GSE68454) and the transcriptome generation by microarray, as well as data handling and analysis were performed as described (Nehar-Belaid et al., 2016).

##### Generation of hypoxia- and vasculogenesis-related gene signatures

Hypoxia (Moslehi et al., 2013) and vasculogenesis (Argraves and Drake, 2005) gene signatures were extracted from the associated publications and their expression were analysed in the total uterine microenvironment (GSE68454) dataset. Then, their power to discriminate the different phenotypic groups was tested by principal component analysis (PCA).

#### Microarray profiling of Tregs coming either from uterus or other tissue

We compared the transcriptome of microarray muscle (MT) (GSE50096), visceral adipose tissue (VAT) (GSE7852) and colonic lamina propria (GSE58164) Tregs with IL18 (GSE71588) and uterine (GSE109895) Tregs. We used the pd.mogene.1.0.st.v1 (MT), pd.mouse430.2 (VAT), illuminaMousev2.db (CLP) R package to annotate the microarray datasets. For RNAseq data, the R package *biomaRt* was rather used. Differentially expressed gene lists from the Tregs of each tissue were obtained comparing the tissue Treg samples with their draining lymph nodes Treg samples for each tissue. DEG analysis was performed for microarray MT, VAT and CLP Tregs with the *limma* package and for IL18 and uterine Tregs with the *DESeq2* package, both using default models and statistical tests (Bayes t-test and Wald test, respectively). DEGs were selected using the following thresholds: false discovery rate < 0.05, |log2FC| ≥ 2 and adjusted p-value < 0.05. The different lists were joined into a table in order to get the expression of all significant genes in every condition.

### TCR Deep Sequencing and Data Processing

Remaining RNAs from NP and E6-E8 uTregs and E6-E8 udLN Tregs used for transcriptome analysis were pooled per group (NP uTregs, E6-E8 uTregs, E6-E8 udLN Tregs). In addition, we obtained Tregs from spleen (NP Spleen), inguinal lymph node (NP Inguinal), pancreatic lymph node (NP Pancreatic) and para-aortic lymph node (NP udLN) from two independent experiments where each sample was a pool of 8 to 10 age-matched C57BL/6 Foxp3^eGFP^ mice. Cells were lysed and RNAs extracted using an RNAqueous kit (Ambion®) as described previously. T cell receptor (TCR) alpha/beta libraries were prepared on 100 ng of RNA from each sample using the SMARTer Mouse TCR α/β Profiling Kit from Takarabio® and sequenced using the MiSeq V2 2×150bp (Illumina®) and HiSeq2500 V2 2×150bp (Illumina®) protocols at the “Institut du Cerveau” (ICM) and at the “LIGAN genomic platform”, respectively (Barennes et al., 2021). Raw data were annotated using MiXCR software (v2.1.10) (Bolotin et al., 2015). MiXCR extracts of TCRs provided corrections of PCR and sequencing errors.

Each dataset is defined as a unique combination of TRV-CDR3aa-TRJ sequences and their associated counts. For NP Spleen, NP Inguinal, NP Pancreatic and NP udLN Tregs obtain from two independent experiment, we pooled clonotypes dataset to obtain unique meta-repertoire comparable in term of mice pooling with NP uTregs, E6-E8 uTregs and E6-E8 udLN Tregs. For Datasets are normalized by a sampling step. The sampling size will be equal to that of the smallest dataset. The sampling is performed for each dataset by 100 random draws of S sequences and unique TRs are listed and counted. After 100 iterations, TRs observed for at least 40 of the 100 are selected and their count is averaged across the number of iterations they were observed in. These values are computed to quantify the differences between repertoires at several complementary levels.

The Generation probability (Pgen) of TRB CDR3aa was inferred using Olga algorithm (Sethna et al., 2019), which is inferred by IGoR (Marcou et al., 2018). Occurrences distributions dendrogram were performed using “PowerTCR” R package (Koch et al., 2018), that provides a model for the clone size distribution of the TCR repertoire. It permits comparative analysis of TCR repertoire libraries based on theoretical model fits. All representations were created using “ggplot2” R package (Hadley Wickham, 2016). A Venn diagram was obtained with the VennDiagram R package (https://cran.r-project.org/web/packages/VennDiagram) .

