## Supplemental Files for "Tissue-resident uterine regulatory T cells support fetal growth"

SUPPLEMENTAL INFORMATION

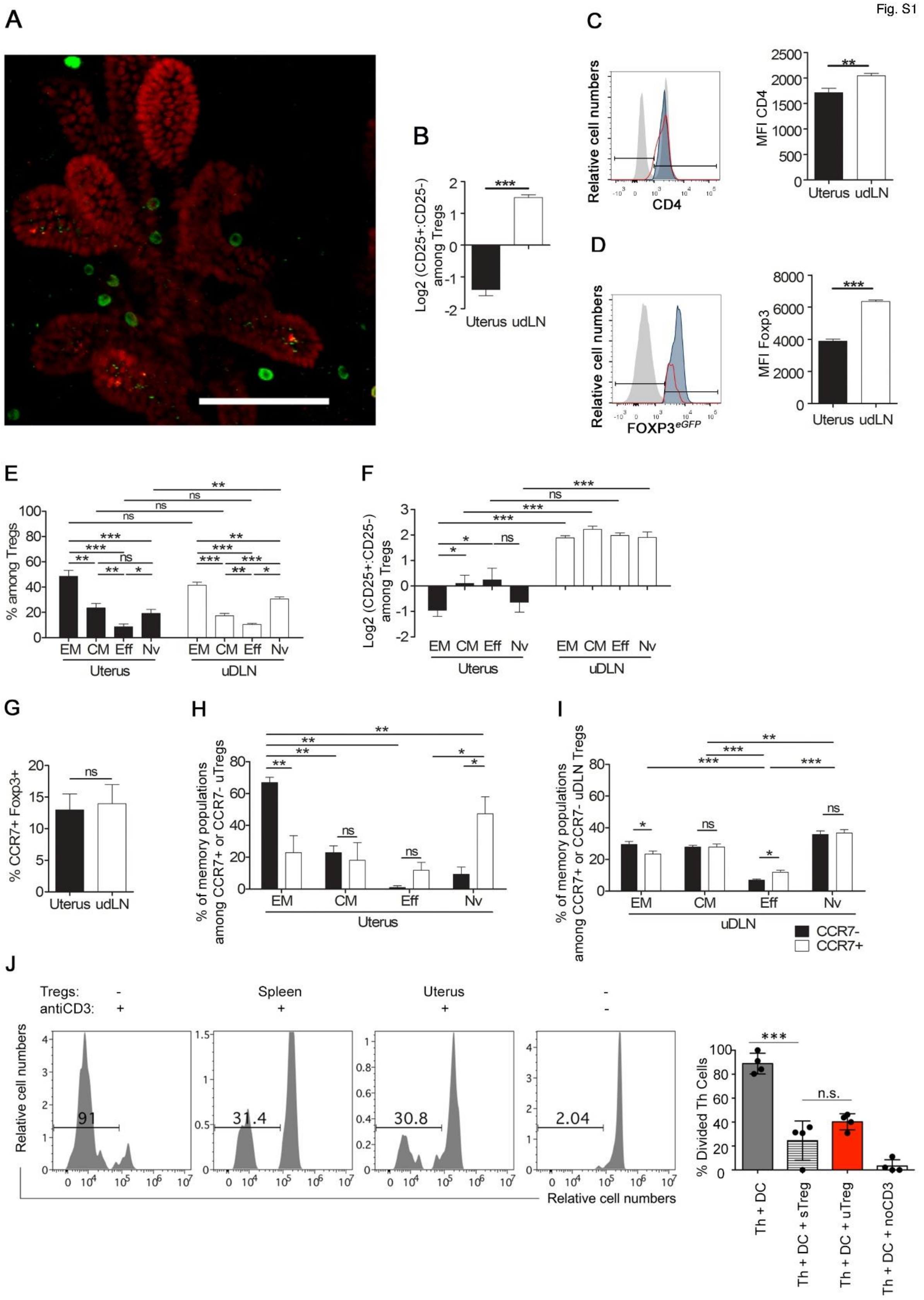
A
B
Log2 (CD25+:CD25-) among Tregs
Uterus udLN
C
Relative cell numbers
CD4
MFI CD4
Uterus udLN
D
Relative cell numbers
FOXP3eGFP
MFI Foxp3
Uterus udLN
E
% among Tregs
EM CM Eff Nv EM CM Eff Nv
Uterus uDLN
F
Log2 (CD25+:CD25-) among Tregs
EM CM Eff Nv EM CM Eff Nv
Uterus uDLN
G
% CCR7+ Foxp3+
Uterus udLN
H
% of memory populations among CCR7+ or CCR7- uTregs
EM CM Eff Nv
Uterus
I
% of memory populations among CCR7+ or CCR7- uDLN Tregs
EM CM Eff Nv
uDLN
CCR7-
CCR7+
J
Tregs: - Spleen Uterus -
antiCD3: + + + -
Relative cell numbers
91
31.4
30.8
2.04
Relative cell numbers
% Divided Th Cells
n.s.
Th + DC
Th + DC + sTreg
Th + DC + uTreg
Th + DC + noCD3

**Supplementary Figure 1. Distribution, phenotypic and residency analysis of uTregs in non-pregnant uteri of Foxp3$^{eGFP}$ mice.**

**A)** In non-pregnant mice, Treg cells localize in the uterine endometrium and gather around the uterine glands. Confocal images of longitudinal sections of *Foxp3$^{eGFP}$* transgenic mouse uteri. Treg cells are identified by immunostaining of GFP (green) and uterine glands by immunostaining of FOXA2 (red). **B)** CD25 expression among uTregs represented as the Log2 of %CD25$^{+}$/%CD25$^{-}$ ratio among Foxp3$^{eGFP+}$CD4$^{+}$ cells (n=12). **(C-D):** Histogram plots show CD4 **(C)** or Foxp3 **(D)** expression by Tregs from uterus (red), uDLNs (blue), and CD4$^{+}$Foxp3$^{eGFP-}$ uterine cells (light gray). Histograms on the right show the expression levels of CD4 (C) or Foxp3$^{eGFP}$ (D) as measured by the mean fluorescence intensity (MFI) among Tregs from Uterus and uterus-draining lymph nodes (udLN).

**E)** Treg naïve and memory populations within the non-perfused uterine tissue. Percentages of effector memory (EM), central memory (CM), effector (Eff) and naïve (Nv) Treg cells based on their expression of CD44 and CD62L. **F)** Log2 of %CD25$^{+}$/%CD25$^{-}$ ratio among the Treg memory populations, (n=5). **(G-I):** Naïve Tregs from peripheral blood express CCR7 while EM uTregs are CCR7$^{-}$. **G)** Histograms show the percentages of CCR7$^{+}$ cells among total uterine and udLN Tregs. **H)** The histograms show the proportions of EM, CM, Nv and Eff subsets among CCR7$^{+}$ Tregs (black) and CCR7$^{-}$ Tregs (red) in the uteri from C57BL/6 naïve females. **I)** The histograms show the proportions of EM, CM, Nv and Eff subsets among CCR7$^{+}$ Tregs (white) and CCR7$^{-}$ Tregs (black) in the udLN from the same mice as I). (n=8, representative of 3 independent experiments). **J)** $VPD_{450}$-labeled CD4$^{+}$Foxp3$^{-}$ helper T cells sorted from the spleen of naive Foxp3$^{eGFP}$ mice were cultured in the presence of dendritic cells and CD3-specific agonist antibody ± Tregs from the spleen or uterus of naive Foxp3$^{gfp}$ mice. $VPD_{450}$ dilution of helper T cells was measured by flow cytometry after 3 days. Numbers indicate percentages of divided cells. Bar graphs on the right summarize data (n=3). *: $p < 0.05$, **: $p < 0.01$, ***: $p < 0.001$, n.s: non-significant.

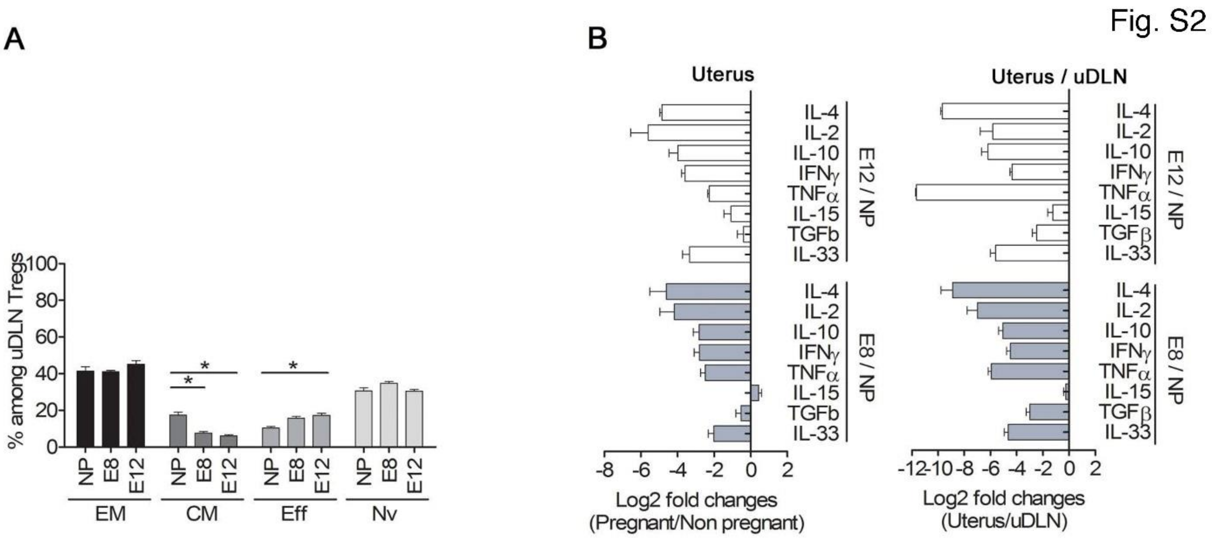


Supplementary Figure 2. Uterine cytokine expression during early pregnancy.

A) udLN Tregs subset is composed of naïve and activated memory cells at E8 and E12. Bar plot representing the percentages of memory populations among udLN Tregs. (n=3 to 5). **B)** T cell-related cytokines are down-regulated in the pregnant uterus. qPCR analysis of T cell-related cytokines. Log2 of fold changes between pregnant and non-pregnant uterus (left panel) and comparison with log2 of fold changes of pregnant uterus with pregnant udLNs (right panel). (n=4).

Fig. S3

| Sample | Organ | Pregnancy | Sequences - TRA | u-TRA | Sequences - TRB | u-TRB |
|---|---|---|---|---|---|---|
| Spleen Treg NP | Superficial LN | Non Pregnant | 42573 | 21142 | 68400 | 38203 |
| Inguinal Treg NP | Superficial LN | Non Pregnant | 44978 | 27606 | 64779 | 41650 |
| Pancreatic Treg NP | Deep LN | Non Pregnant | 42593 | 12604 | 76530 | 25238 |
| dLN Treg NP (para-aortic) | Deep LN | Non Pregnant | 41696 | 12765 | 77349 | 27862 |
| dLN Treg E6-E8 (para-aortic) | Deep LN | Pregnant | 45504 | 29904 | 67438 | 49768 |
| uTregs NP | Uterus | Non Pregnant | 39311 | 6356 | 70283 | 9238 |
| uTregs E6-E8 | Uterus | Pregnant | 39033 | 1447 | 70118 | 2719 |

Supplementary Figure 3. TCR repertoire of uTregs.

Sample properties: Sample: Organ – cell_subset – pregnancy state; Organ: Organ category; Pregnancy: pregnancy state; Sequences – TRA: number of productive TCR alpha chain sequences; u-TRA: number of unique amino acid TCR alpha clonotypes; Sequences – TRB: number of productive TCR beta chain sequences; u-TRB: number of unique amino acid TCR beta clonotypes.

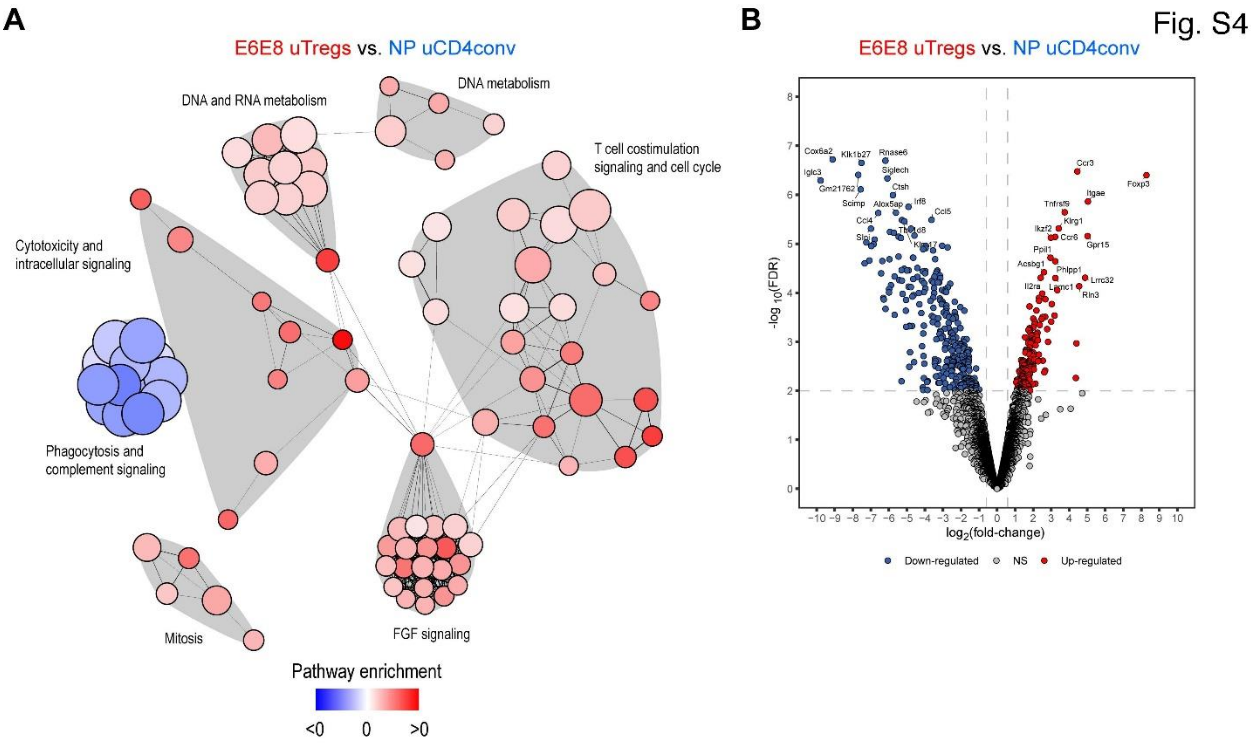


### Supplementary Figure 4. RNA-Seq analysis of uTregs

A) Network representation of pathway enrichment analysis from the list of pathways statistically differentially expressed between E6-E8 uTregs and NP uCD4conv. Each node in the representation corresponds to a pathway, and nodes are connected if the lists of genes in the associated pathways are overlapping. Nodes are colored using a color-gradient scale ranging from blue (<0) to red (>0) according to the difference of mean expressions between the conditions. Clusters of pathways are distinguished from each other by their grey-colored area. B) Volcano plot showing genes differentially expressed between uTregs E6-E8 and NP uCD4conv.

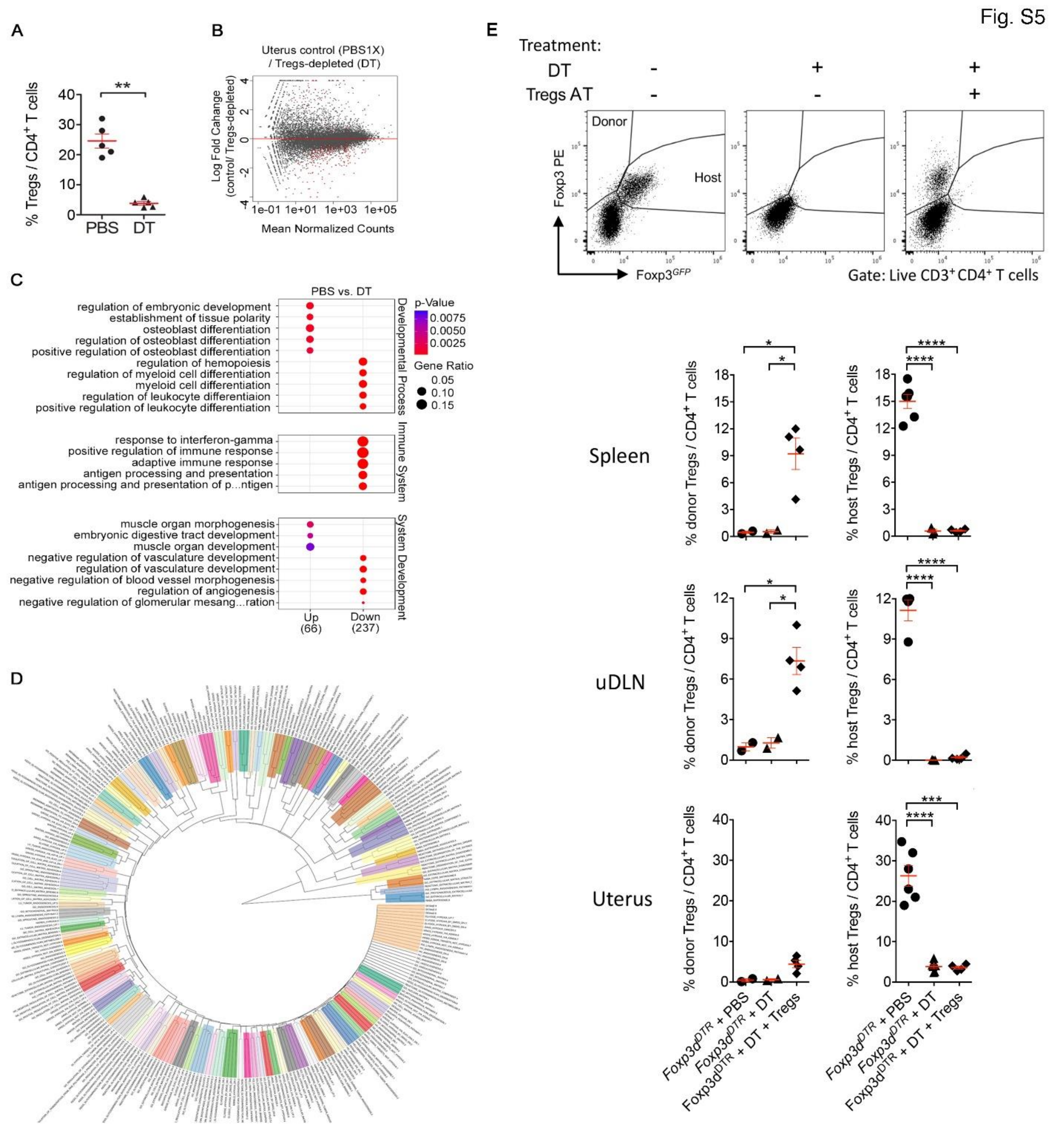


### Supplementary Figure 5. uTregs biological function during pregnancy

**A)** Percentage of Tregs among CD4 total uterine cells comparing Tregs-depleted animals (DT) with controls (PBS), from pregnant E6-E8 females Foxp3$^{DTR}$, used for transcriptome experiments. (**=p-value=0.0079). RNA-Seq analysis from Foxp3$^{DTR}$ total uterus control (PBS1X) vs. Tregs-depleted

(DT) (n=4). **B)** MA-plot shows RNA transcript fold change as a function of transcript abundance (mean normalized counts) between the comparisons: control (PBS1X) vs. Treg-depleted (DT). Differentially expressed genes (false discovery rate (FDR) = 5%; | $\log_2$ normalized expression | > 2) are in red. High abundance transcripts appear on the right x-axis. Common genes are mostly shown in black. **C)** Differential transcriptional analysis of total uteri after Tregs depletion in the $Foxp3^{DTR}$ mouse line. Results of gene ontology (GO) enrichment analysis to examine biological processes of genes differentially expressed between control (PBS) vs. the Tregs-depleted (DT) total uterus transcriptome from $Foxp3^{DTR}$ mouse. The p-value (< 0.01) is represented by colors and the gene ratio by circle sizes. **D)** Binary distance tree used to create 'Modules' (Figure 7D) by cutting neighboring clusters of all signatures from two gene set collections, the C2 and the C5 from the Molecular Signatures Database (MSigDB), for the following terms *angiogenesis*, *vasculogenesis*, *vascularization*, *extracellular matrix*, *basement membrane*, *glycosaminoglycans*, *hyaluronic acid*, *heparan sulfate*, *perlecan*, *collagen*, *elastin*, and *laminin*. **E)** Representative dot plots (top) and histograms (bottom) show the proportions of donor ($Foxp3\text{-}PE^{+}$ $Foxp3^{GFP-}$) and host ($Foxp3\text{-}PE^{+}Foxp3^{GFP+}$) Tregs among $CD3^{+}CD4^{+}$ total cells at E8 in the indicated organs from control PBS- or DT- treated $Foxp3^{DTR}$ mouse, or DT- treated $Foxp3^{DTR}$ mice that received $9 \times 10^6$ Tregs cells from WT mice at E1.5 (n = 2 to 7 mice per group). A Student's t-test was used for statistical analysis. *: P < 0.05; **: P < 0.01; ***: P < 0.001; ****: P < 0.0001. All significant statistics are displayed.
